# MalKinID: A Likelihood-Based Model for Identifying Malaria Parasite Genealogical Relationships Using Identity-by-Descent

**DOI:** 10.1101/2024.07.12.603328

**Authors:** Wesley Wong, Lea Wang, Stephen S Schaffner, Xue Li, Ian Cheeseman, Timothy J.C. Anderson, Ashley Vaughan, Michael Ferdig, Sarah K Volkman, Daniel L Hartl, Dyann F Wirth

## Abstract

Pathogen genomics is a powerful tool for tracking infectious disease transmission. In malaria, identity-by-descent (IBD) is used to assess the genetic relatedness between parasites and has been used to study transmission and importation. In theory, IBD can be used to distinguish genealogical relationships to reconstruct transmission history or identify parasites for genotype-to-phenotype quantitative-trait-locus experiments. *MalKinID* (Malaria Kinship Identifier) is a new likelihood-based classification model designed to identify genealogical relationships among malaria parasites based on genome-wide IBD proportions and IBD segment distributions.

*MalKinID* was calibrated to the genomic data from three laboratory-based genetic crosses (yielding 440 parent-child and 9060 full-sibling comparisons). *MalKinID* identified lab generated F1 progeny with >80% sensitivity and showed that 0.39 (95% CI 0.28, 0.49) of the second-generation progeny of a NF54 and NHP4026 cross were F1s and 0.56 (0.45, 0.67) were backcrosses of an F1 with the parental NF54 strain. In simulated outcrossed importations, *MalKinID* accurately reconstructs genealogy history with high precision and sensitivity, with F1- scores exceeding 0.84. However, when importation involves inbreeding, such as during serial co-transmission, the precision and sensitivity of *MalKinID* declined, with F1-scores of 0.76 (0.56, 0.92) and 0.23 (0.0, 0.4) for PC and FS and <0.05 for second-degree and third-degree relatives. Genealogical inference is most powered 1) when outcrossing is the norm or 2) when multi-sample comparisons based on a predefined pedigree are used. *MalKinID* lays the foundations for using IBD to track parasite transmission history and for separating progeny for quantitative-trait-locus experiments.

## Introduction

*Plasmodium falciparum* is a single-cell, eukaryotic parasite that is responsible for one of the deadliest forms of malaria. Global eradication efforts have drastically altered the transmission landscape of *P. falciparum* and malaria parasite genomics has emerged as a powerful tool for assessing ongoing transmission and the effectiveness of public health interventions (Ashton *et al*. 2020; Wong *et al*. 2023). However, the population genomics of the malaria parasite is unusual among pathogens because it must sexually reproduce and recombine in a mosquito vector during transmission. As such, each parasite strain reflects up to two different genealogical histories, one from each parent, and can be genetically related to other individuals in the population.

The genetic relatedness of malaria parasites is a robust metric for malaria genomic epidemiology studies and has been used to evaluate changes in transmission (Neafsey and Volkman 2017; Taylor *et al*. 2017; Neafsey *et al*. 2021), distinguish between hypotheses (superinfection vs. co-transmission) of multiple strain (polygenomic) infection formation (Nkhoma *et al*. 2012, 2018, 2020; Nair *et al*. 2014; Wong *et al*. 2017, 2018, 2022), detect differences in transmission structure (Schaffner *et al*. 2023), and identify regions of the genome undergoing strong, directional selection (Schaffner *et al*. 2023). In these studies, genetic relatedness is defined as the proportion of the genome between two individuals that shows identity-by-descent (IBD) and thus inherited from the same recent common ancestor (**Figure 1**).

**Figure 1.**
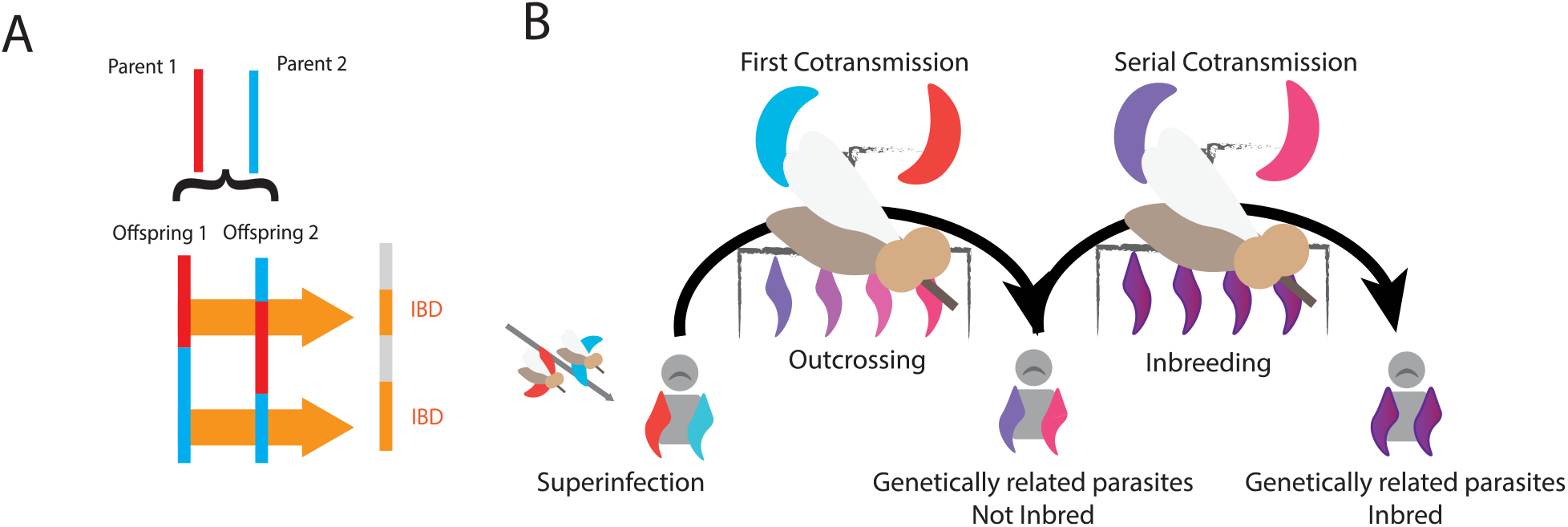
**A**) Definition of genetic relatedness as the proportion of the genome that is considered Identity-by-Descent (IBD). IBD is defined as a section of a genome that is shared between two individuals and inherited from the same recent common ancestor. **B**) Generation of genetically related malaria parasites. Genetically related parasites first appear when a mosquito bites a superinfected individual with unrelated parasite strains. This provides parasites their first opportunity to outcross (mate between a genetically unrelated strain) and produce genetically related, but not inbred, progeny that are then transmitted to the next individual. If multiple, genetically distinct but related parasites are transmitted, the transmission event is referred to as co-transmission. Subsequent transmission and serial co-transmission events provide opportunities for inbreeding (mating between genetically related strains) and can generate genetically related, inbred progeny.

Despite its use in malaria genomic epidemiology, parasite genetic relatedness can be difficult to interpret. Partially related parasites are the result of a complex series of transmission events that involve a mix of both superinfection and co-transmission (**Figure 1B**). Superinfection occurs when an individual is infected with multiple strains from separate mosquito bites. The strains within this initial superinfection represent a random sampling of parasites from the population and are assumed to be unrelated (Nkhoma *et al*. 2012, 2020; Wong *et al*. 2017, 2022).

Mosquitoes feeding on this superinfected polygenomic infection provide the parasite with its first opportunity to outcross and generate genetically related progeny. Co-transmission occurs when the mosquito feeds on the superinfected infection and transmits multiple, often genetically related parasites into a new host. Subsequent feedings on these co-transmitted polygenomic infections enable inbreeding and the generation of inbred progeny. As such, the genetic relatedness of parasites depends on transmission history and how superinfection and co-transmission occur in the population.

New approaches that can disentangle the complex relationship between transmission history and genetic relatedness could be valuable for future malaria genomic epidemiology studies. In theory, transmission lineages could be reconstructed by identifying parent-child relatives in the population. Genealogical inference has been used in conservation biology and ecology to reconstruct pedigrees and study recent demographic history or monitor wild animal population movement (Blouin 2003; Kirkpatrick *et al*. 2011; Staples *et al*. 2014; Huisman 2017; Jones and Manseau 2022). Genealogical inference in malaria would also be useful for establishing inbred parasites lines for laboratory-based quantitative-trait-locus (QTL) experiments (Daley and Shepherd 2008; Solberg Woods 2014) designed to identify molecular determinants of antimalarial drug resistance.

However, genealogical inference is a surprisingly nuanced problem in malaria. One consequence of its haploid genome is that progeny parasites are not guaranteed to inherit half of their genomes from each parent (Wong *et al*. 2018). Another consequence is that genetically distinct meiotic siblings from the same oocyst, a sac-like structure that forms in the mosquito midgut after zygosis and contains the immediate, haploid products of meiosis, have an expected relatedness of 0.33 (Wong *et al*. 2018; Nkhoma *et al*. 2020). Meiotic siblings are distinct from full siblings, which have an expected relatedness of 0.5, and occur when two, genetically distinct haploid parasites are sampled from two different oocysts that involve the zygosis and meiosis of the same parental strains (Wong *et al*. 2018; Nkhoma *et al*. 2020).

*MalKinID* (*Malaria Kinship Identifier*) is a new likelihood-based classification model that infers the genealogical relationship between parasite strains based on the genome-wide IBD proportion and the per-chromosome maximum IBD segment block (*IBDmax*) and IBD segment count (*nsegment*) distributions. The likelihoods used in *MalKinID* were based on the patterns observed in simulated parasites generated using a previously developed meiosis model with obligate chiasma formation and crossover interference (Wong *et al*. 2018) that was calibrated to whole genome sequencing data obtained from three laboratory-based genetic crosses [NF54 × NHP4026, MKK2835 × NHP1337, and MAL31 × KH004] (**Figure 2**).

**Figure 2.**
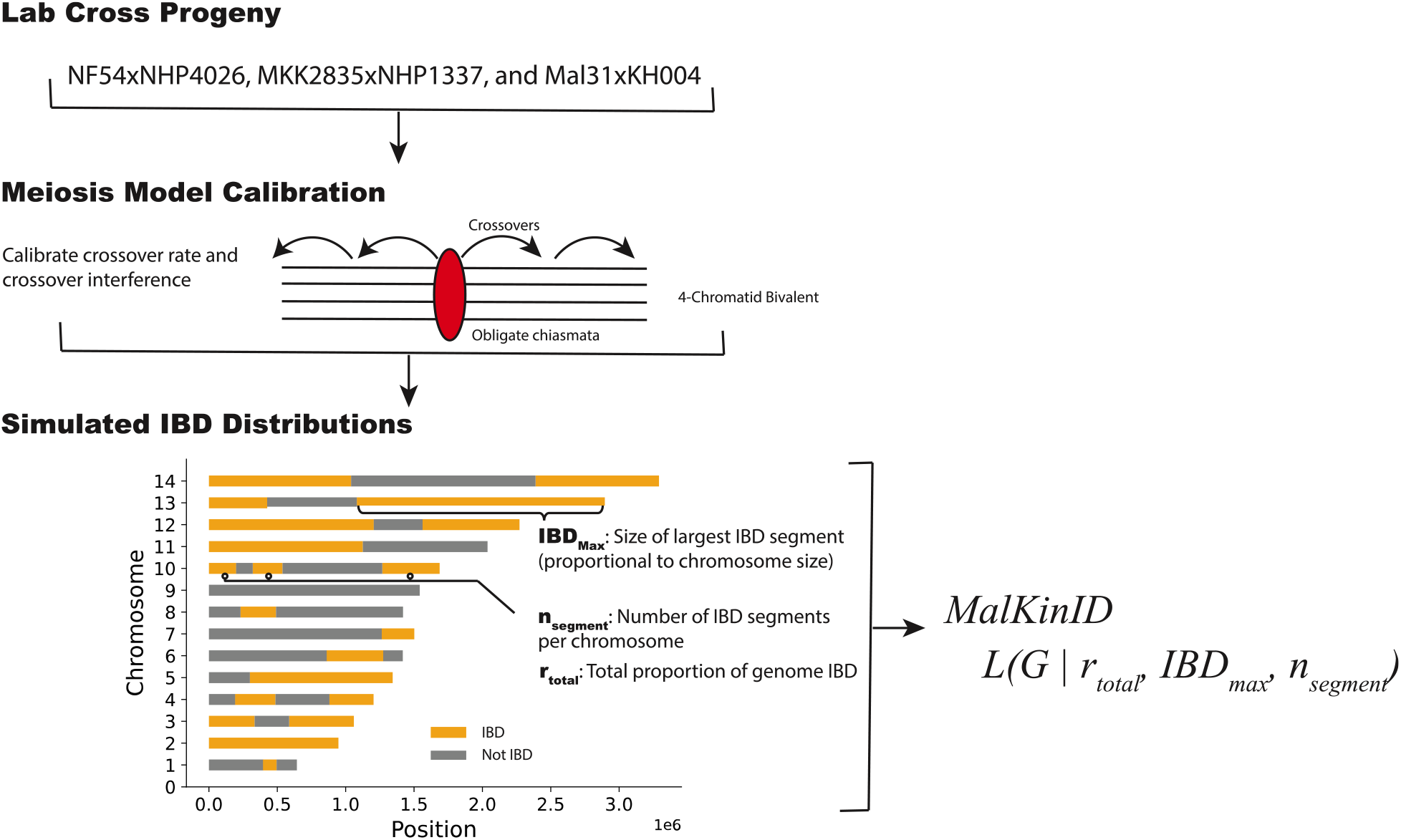
Study Design. *MalKinID* infers the genealogical relationship (G) between two parasites based on *r*_*total*_, *IBD*_*max*_, and *n*_*segments*_. The likelihood is based on using a calibrated meiosis model that includes obligate chiasma formation and crossover interference to simulate the distributions for *r*_*total*_, *IBD*_*max*_, and *n*_*segments*_ for each genealogical relationship.

## Results

### Brief model overview

By default, *MalKinID* classifies parasite pairs into eleven different genealogical relationships (**Table 1**) that cover a broad range of outcrossed parasite relationships. These genealogical relationships include first-degree relatives (parent-child [PC], full-sibling [FS], and meiotic siblings [MS]), second-degree relatives (grandparent-grandchild [GC], half-siblings [HS], full avuncular relationships where the avuncular relative is a FS [FAV] or MS [FAV.MS] to the parent), and third-degree relatives (great-grandparent – great-grandchild [GGC], full cousin where the related parental strains are FS [FCS] or MS [FCS.MS], and half-avuncular [HAV]). *MalKinID* is not restricted to these genealogical relationships and can be modified to perform multi-sample comparisons to identify any genealogical relationships based on a pre-defined pedigree tree.

**Table 1.** The genealogical relationships examined in this study and their expected genetic relatedness values. Red highlighting is used for first-degree relatives, blue for second-degree relatives, grey for third-degree relatives, and white for unrelated pairs. Bold and italicized rows emphasize the relationships that represent vertical genealogical descent.

| Kinship | Degree | Expected Genetic Relatedness |
| --- | --- | --- |
| ***Parent-Child (PC)*** | ***First*** | ***0.5*** |
| Full-Siblings (FS) | First | 0.5 |
| Meiotic Sibling (MS) | First | 0.33 |
| ***Grandparent-Grandchild (GC)*** | ***Second*** | ***0.25*** |
| Half Siblings (HS) | Second | 0.25 |
| Full Avuncular (FAV and FAV.MS) | Second | 0.25 / 0.166 |
| ***Great Grandparent-Great Grandchild (GGC)*** | ***Third*** | ***0.125*** |
| Half Avuncular (HAV) | Third | 0.125 |
| Full Cousins (FCS and FCS.MS) | Third | 0.125 / 0.0830 |
| Unrelated |  | 0 |

### Simulating the genetic relatedness of parasite pairs using a calibrated meiosis model

Previous pedigree reconstruction studies have shown that the genome-wide proportion of IBD between two individuals (referred to as total relatedness or *r*_*total*_) and the IBD segment block distribution can be used to distinguish genealogical relationships (Browning 1998; Zhao and Liang 2001). To test whether these findings applied to malaria, a previously published meiosis model that includes obligate chiasmata formation and crossover interference (Wong *et al*. 2018) was used to derive the distributions for *r*_*total*_ and the per-chromosome (*IBD*_*max*_, proportional to the size of each chromosome) and IBD segment count (*n*_*segment*_) for each of the genealogical relationships described in **Table 1** based on the pedigree described in **Figure 3A** (**Methods, Equations 1-4**). The distribution of IBD segment blocks was summarized by using the per-chromosome values of *IBD*_*max*_ and *n*_*segment*_. These values were chosen because they provide statistically independent measures for each chromosome, which facilitates their integration into the final likelihood model.

**Figure 3.**
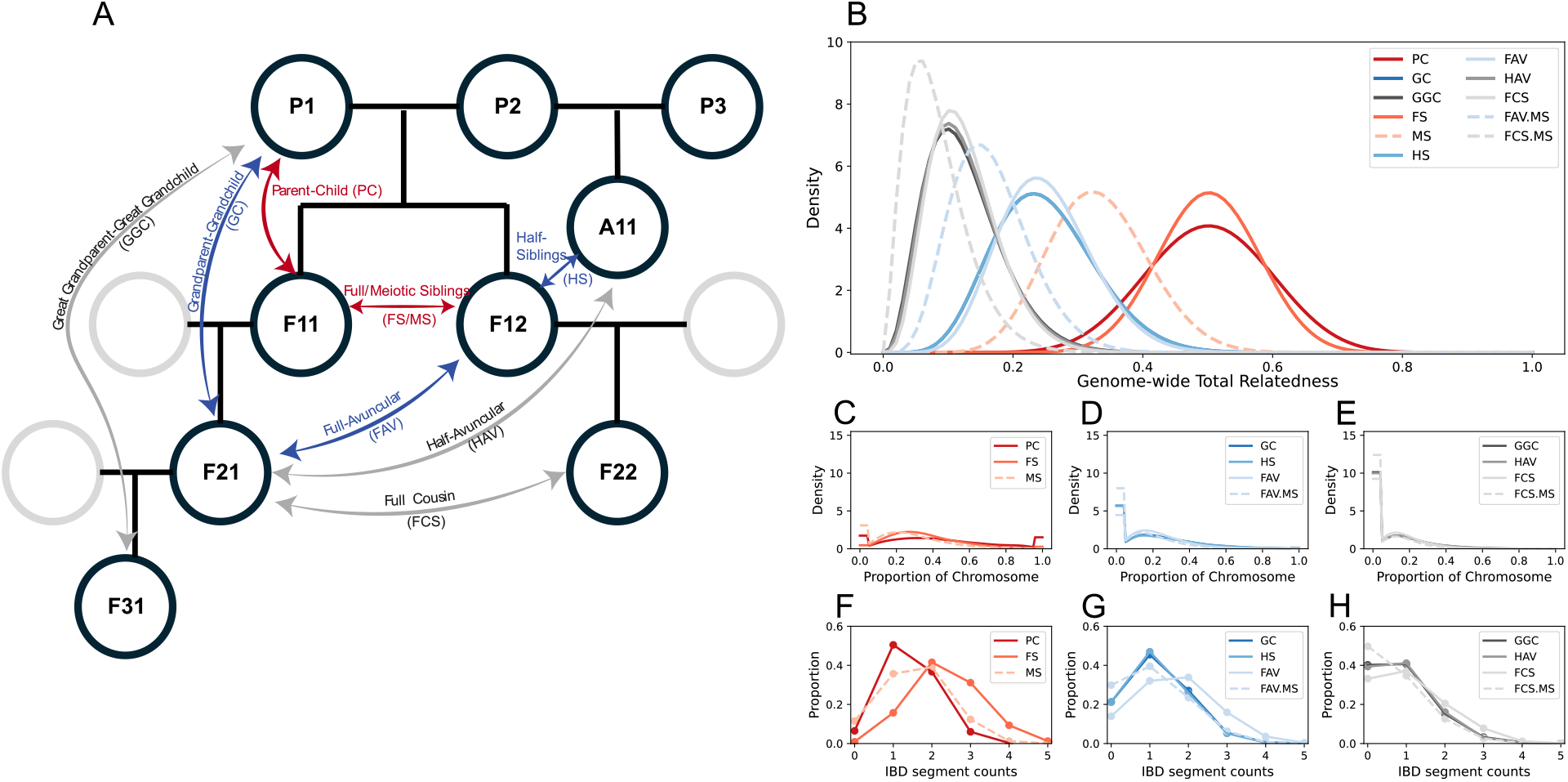
Simulating outcrossed pedigrees **A**) The pedigree used to simulate genetically related parasites in this study. Each node of the pedigree represents a genetically distinct parasite. Unlabeled notes represent external, genetically unrelated parasites that are used for outcrossing. The parental strains (P1, P2, and P3) are all genetically unrelated to one another. F11 and F12 represent two different F1-progeny, F21 and F22 two different F2-progeny, and F31 an F3 progeny descended from P1 and P2. A11 refers to a descendent from an “alternate” lineage involving P2 and P3. Simulated distribution for **B**) genome-wide total relatedness (*r*_*total*_), max IBD segment (*IBD*_*max*_) distributions for chromosome 14 for **C**) first-, **D**) second-, and **E**) third-degree relatives, and IBD segment count (*n*_*segments*_) distribution for chromosome 14 for **F**) first-, **G**) second-, and **H**) third-degree relatives. These simulated distributions were generated by fitting the raw simulation output to the equations used in the likelihood model (**Equations 1-4**). FAV.MS and FCS.MS denote FAV and FCS descended from MS.

The meiosis model incorporates obligate chiasma formation and includes parameters that specify the crossover rate and strength crossover interference. For this study, the meiosis model was recalibrated to newly available genomic data obtained from 440 unique PC relationships and 9060 unique FS relationships from three laboratory-crosses: NF54 x NHP4026, MKK2835 x NHP1337, and Mal31 x KH004 (**Supplemental Table 1, Supplemental Figure 1, Supplement Text**). Several optimum parameter choices were observed (**Supplemental Figure 1**): 1) a set with no crossover interference, 2) a set with weak crossover interference, and 3) a set with moderate/high crossover interference (see **Supplemental Text:** *Meiosis model recalibration* for additional details). The parameter set with no crossover interference was excluded because crossover interference is a fundamental aspect of meiosis that is observed across diverse taxa. Unless otherwise noted, all model results were made using the weak crossover interference parameter set because it results in more accurate PC classification rates (see below and **Supplemental Text**).

*Simulations reveal variations in genetic relatedness and IBD segment block distribution between genealogical relationships*.

These simulations revealed differences in *r*_*total*_ and the IBD segment block distributions that could be used to distinguish genealogical relationships (**Figure 3**, **Supplemental Figure 2**). First-degree relatives were the most related, with expected *r*_*total*_s of 0.50 for PC and FS and 0.33 for MS. Simulations showed that the IBD segment block distributions were significantly different for each of these genealogical relationships (p-value << 1.0e-10 for all chromosomes (**Figure 3F**, Kruskal-Wallis Test).

Relative to FS, PC have fewer but larger IBD segments that were more likely to span the entire length of the chromosome. For PC, smaller chromosomes, such as chromosome 1, were more likely to have chromosome wide IBD segments (0.23, [95%CI 0.22, 0,24]) than longer chromosomes (*e.g.* chromosome 14, 0.07 [95% CI 0.06, 0.07]) (**Supplemental File 3**). These differences in the IBD segment block distribution explain why the distribution of *r*_*total*_ for FS was narrower than that of PC, despite having identical expectations (**Supplemental Figure 3**). The standard deviation of the *r*_*total*_ distributions from the simulated PC and FS distributions were 0.095 (95% CI 0.093, 0.097) and 0.076 (95% CI 0.075, 0.078), respectively.

With the exception of full avuncular and full cousin relationships, the expected relatedness for all second- and third-degree relatives were 0.25 and 0.125, respectively (**Figure 3B**). The expected relatedness of full avuncular relatives and full cousins depended on whether their parental F1 strains (nodes F11 and F12 in **Figure 3A**) were FS or MS. If the parental F1 strains were FS (FAV and FCS), they had the same expected *r*_*total*_ as the other second- and third-degree progeny, 0.25 and 0.125 respectively. If the parental F1 strains were MS (FAV.MS and FCS.MS), they were more unrelated than expected, with an expected *r*_*total*_ of 0.166 and 0.0830, respectively.

Overall, the IBD segment block distributions of second- and third-degree relatives were characterized by fewer IBD segments and a greater proportion of chromosomes with no IBD segments at all (**Figure 3 D-E, G-H**). While each of the genealogical relationships within second- and third-degree relatives had significantly different *IBD*_*max*_ and *n*_*segment*_ distributions (p-value << 1.0e-10 for all chromosomes, Kruskal-Wallis Test), the differences were less pronounced than those seen within first-degree relatives.

*Accurate genealogical inference required utilizing* *r*_*total*_ *and the per-chromosome* *IBD*_*max*_*, and* *n*_*segment*_ *distributions*

Based on these results, the differences in *r*_*total*_ and the per-chromosome *IBD*_*max*_ and *n*_*segment*_ distributions were evaluated to determine whether they could be used for genealogical inference. It was hypothesized that a likelihood model utilizing *r*_*total*_ alone could distinguish first-, second-, and third-degree relatives but that further classification into the specific genealogical sub-relationships (for instance, PC and FS) would require a likelihood model that also utilized the per-chromosome *IBD*_*max*_and *n*_*segment*_distributions. It was also of interest to determine whether including MS, FAV.MS, and FCS.MS would lessen genealogical inference accuracy, as their respective *r*_*total*_ distributions blurred the distinctions between the other first-, second-, and third-degree relatives (**Figure 3B**). To address these, this study evaluated several sets of likelihoods that involved different modifications of the likelihood described in **Equation 1**.

The first set of examined likelihoods excluded the MS, FAV.MS, and FCS.MS categories and focused on evaluating whether a likelihood model utilizing *r*_*total*_ alone could accurately classify genealogical relationships. As expected, classification based on *r*_*total*_ alone performed poorly when identifying individual genealogical relationships but could be used to classify parasite pairs as first-, second-, or third-degree relatives (**Supplemental Figure 4**). The true classification rate (the proportion of accurately identified comparisons) for first-degree relatives was 0.943 (95% CI 0.938, 0.947), second-degree relatives 0.771 (95% CI 0.762, 0.779) and third-degree relatives 0.828 (95% CI 0.821, 0.836).

Including the per-chromosome *IBD*_*max*_and *n*_*segment*_distributions into the likelihood model increased the true classification rates for first- 0.989 (95% CI 0.987, 0.991), second- 0.925 (95% CI 0.92, 0.93), and third-degree relatives 0.944 (95% CI 0.940, 0.950) (**Supplemental Figure 5**). It also allowed the model to further distinguish first-degree relatives as either PC (0.96 [95% CI 0.956, 0.967]) or FS (0.97 [95% CI 0.964,0.974]). However, differentiation of the genealogical relationships within second- and third-degree relatives remained challenging, with true classification rates of ∼0.40 for GC, HS, GGC, and HAV and 0.6 – 0.80 for FAV and FCS.

These trends held true when expanding the likelihood model to include meiotic siblings and their descendants (**Figure 4A-E**). Overall, including MS, FAV.MS, and FCS.MS caused true classification rates to decline slightly, particularly for the genealogical relationships that define second- and third-degree relatives. As before, *r*_*total*_ alone was insufficient to accurately distinguish individual genealogical relationships. When including the additional MS relationships in the model, the true classification rates for first-, second-, and third-degree relatives was 0.93 (95% CI 0.92, 0.94), 0.81 (95% CI 0.80, 0.83), and 0.65 (95% CI 0.64, 0.66). The classification rates for the individual genealogical relationships within first-degree relatives were consistently higher than those for second- and third-degree relatives. The true classification rate for MS was 0.76 (95% CI 0.75, 0.77).

**Figure 4.**
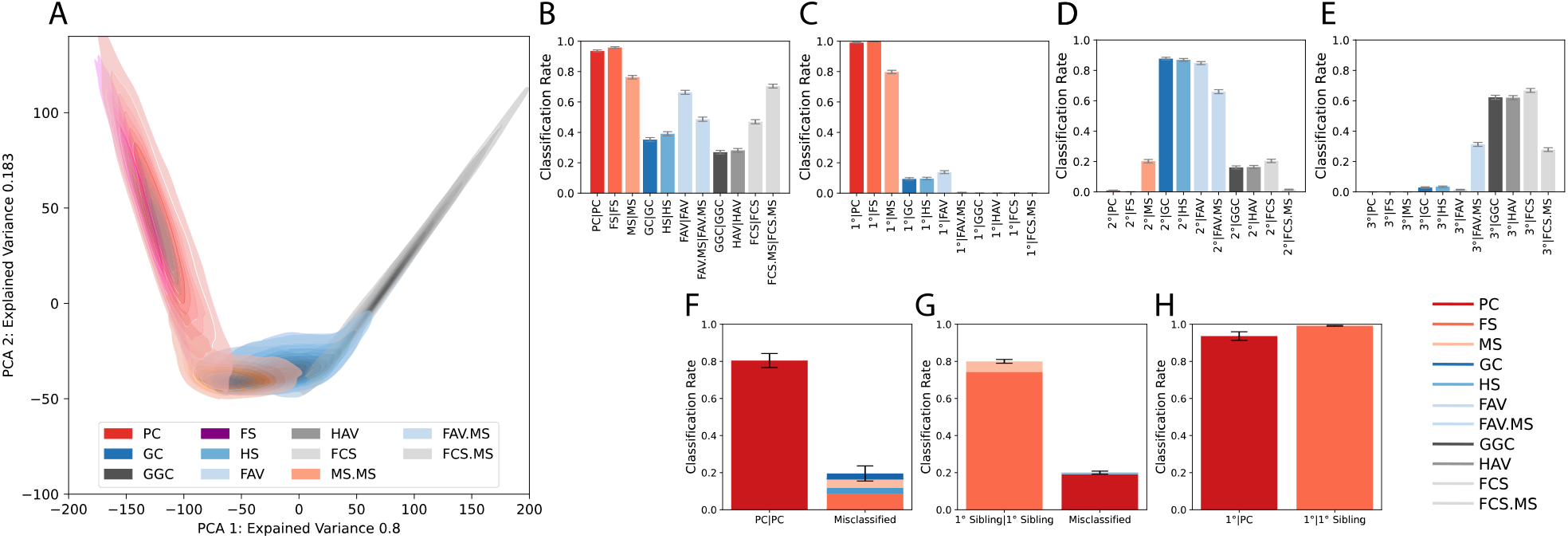
Classification of outcrossed parasite genealogies on simulated (**A-E**) and empirical lab-cross data (**F-H**). Refer to Figure 2A or **Table 1** for abbreviations. A) PCA plot of the simulated *r*_*total*_, *IBD*_*max*_ and *n*_*segment*_ data. The PCA included a third component that is not shown. Each genealogical relationship is represented by 5000 independent simulations and was fit with a gaussian kernel density estimator for visual clarity. Darker spots indicate a region with higher mass for that genealogical relationship. *MalKinID* classification rates when including all meiotic sibling possibilities (**B-E**). The term to the left of the “|” refers to the inferred classification and the term to the right the true genealogical relationship. **B** classifies samples by their genealogical relationship while the other subplots classify samples as (**C**) first-, (**D**) second-, or (**E**) third-degree relatives. MS.MS refers to meiotic siblings, while FAV.MS and FCS.MS refer to the meiotic sibling variants for FAV and FCS. Empirical classification rates for (**F**) PC, (**G**) first- degree siblings (FS and MS), and (**H**) first-degree relatives from the laboratory cross data. Error bars indicate two standard deviations from the mean from 5000 comparisons. 1^0^ siblings refers to FS and MS. 1^0^ refers to all first-degree relatives (PC, FS, and MS).

### MalKinID accurately classifies known PC and FS relatives from three laboratory-based genetic crosses

The performance of *MalKinID* was first assessed using the genomic data from the three crosses (NF54 x NHP4026, MKK2835 x NHP1337, Mal31 x KH004) (**Fig 4F-H**). In these crosses, *MalKinID* correctly identified 0.80 (95% CI 0.77, 0.84) of the PC relationships and 0.80 (95% CI 0.79, 0.81) of the first-degree siblings (FS or MS). Of the correctly identified first-degree siblings, *MalKinID* 0.92 (95% CI 0.91, 0.94) were identified as FS. The average of the true classification rates for PC and first-degree siblings (the macro-average) was 0.80 (95% CI 0.77, 0.83). When using *MalKinID* to classify parasites as first-, second- or third-degree relatives, 0.94 (95% CI 0.91, 0.96) and 0.99 (95% CI 0.98, 0.99) of the PC and FS relationships were correctly identified as first-degree relatives and the overall, macro-average rate for both PC and FS comparisons was of 0.96 (95% CI 0.93, 0.99).

Replacing the *r*_*total*_, *IBD*_*max*_, and *n*_*segments*_ distributions with those generated from the meiosis model with the moderate/strong crossover interference parameter set (**Supplemental Text**) resulted in similar macro-average classification rates across all PC and FS comparisons [0.80 (95% CI 0.76,0.84)], but differential true classification rates for FS and PC. This parameterization yielded higher true classification rates for FS [0.92 (95% CI 0.91, 0.93)] but worse true classification rates for PS [0.68, (95% CI 0.63, 0.72)] (**Supplemental Figure 6, Supplemental Text**).

### MalKinID accurately reconstructs the genealogical history of outcrossed parasite lineages with high precision and recall

To determine whether *MalKinID* could faithfully reconstruct transmission lineages, the performance of *MalKinID* was evaluated using simulations designed to represent: 1) “outcrossed point importations”, the importation of a single parasite strain in a population with sufficiently high transmission intensity that superinfection (repeated infection of an already infected individual) promotes parasite outcrossing (**Figure 5A/C**), and 2) “inbred point importations” (**Figure 5B/D**), the importation of a single parasite strain in a population with such low transmission intensity or such isolated transmission lineages that superinfection is rare and serial co-transmission chains (the repeated transmission of multiple, often genetically-related parasites) are the norm. For outcrossed point importations, *MalKinID* accurately identified parasite relationships with high precision and sensitivity (**Figure 5C**). The F1-scores for PC, FS, second-degree, and third-degree relatives were 0.95 (0.84, 1.0), 0.94 (0.72, 1.0), 0.84 (0.60, 1.0) and 0.84 (0.64, 0.94), respectively.

**Figure 5.**
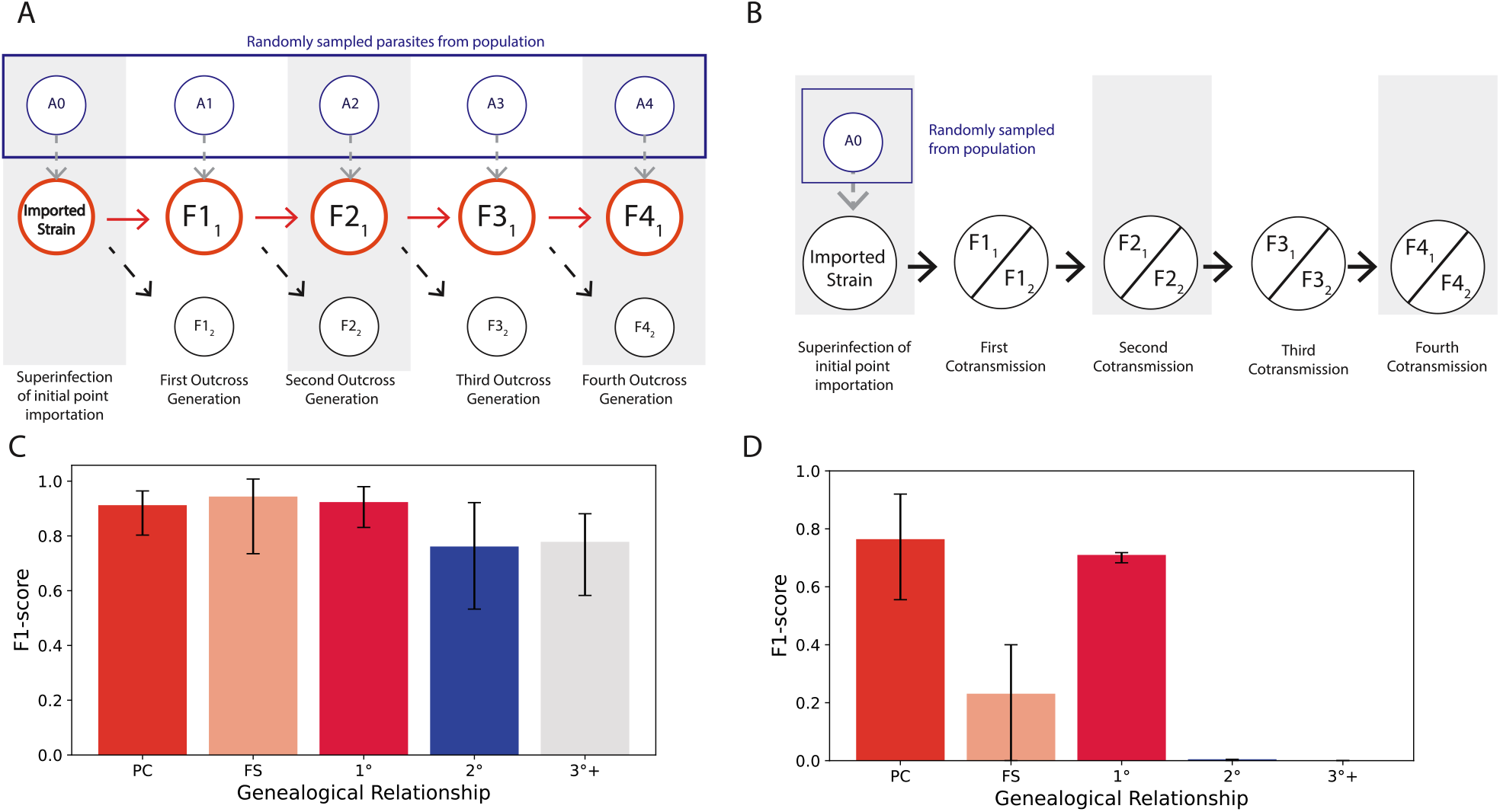
Performance of *MalKinID* on simulated importations involving **A/C**) outcrossing and **B/D**) inbreeding through serial co-transmission. In **A** and **B**, nodes starting with “A” represent superinfecting parasites sampled from the local population. For all other nodes, the non- subscript part of the names denotes the generation (F1, F2, F3, and F4) while the subscript refers to different versions of it. The accuracy of *MalKinID* was summarized using the F1-score. The error bars indicate two standard deviations from the mean.

However, there was a significant drop in precision and sensitivity when *MalKinID* was applied to inbred point importations, with F1-scores of 0.76 (0.56, 0.92) and 0.23 (0.0, 0.4) for PC and FS and <0.05 for second-degree and third-degree relatives (**Figure 5D**). Under these conditions, second- and third-degree relatives were consistently misidentified as PC. This is because inbreeding can result in a wide range of overlapping *r*_*total*_ and per-chromosome *IBD*_*max*_ and *n*_*segment*_ distributions whose values depend on how inbred the original parental strains are (**Supplemental Figure 7**).

### Tree-based genealogical inference and multi-sample comparisons can disentangle inbred parasite genealogies

Thus far, the analyses presented in this study have focused on using *MalKinID* to identify the most likely genealogical relationship between parasite pairs. **Figure 5D** showed that relying on pairwise comparisons to perform genealogical inference fails when inbreeding occurs.

Fortunately, the issues caused by inbreeding can be overcome if inferences using multi-sample comparisons (Sieberts *et al*. 2002) based on a pre-defined pedigree that describes all the possible genealogical relationships between sample pairs are used. Such pedigrees are difficult to specify in genomic epidemiology studies but are readily available in laboratory-based genetic crosses and QTL experiments.

As an example, *MalKinID* was used to identify outbred and inbred parasites from a two-generation cross involving NF54 and NHP4026 (**Figure 6A**). This two-generation cross was generated by feeding mosquitoes a blood meal containing NF54 and NHP4026, pooling the resulting first-generation progeny, and feeding that pool to a second batch of mosquitoes. The initial pool of first-generation progeny contained a mix of F1 progeny generated through outcrossing (mating between two genetically distinct parasites) and parental clones generated through selfing (mating between two clones). This pool is then fed to a second mosquito without prior removal of parental clones. As a result, the second-generation pool could contain parental, F1, and F2 progeny as well as backcrossed progenies generated by crossing an F1 with NF54 (B1) or NHP4026 (B2).

**Figure 6.**
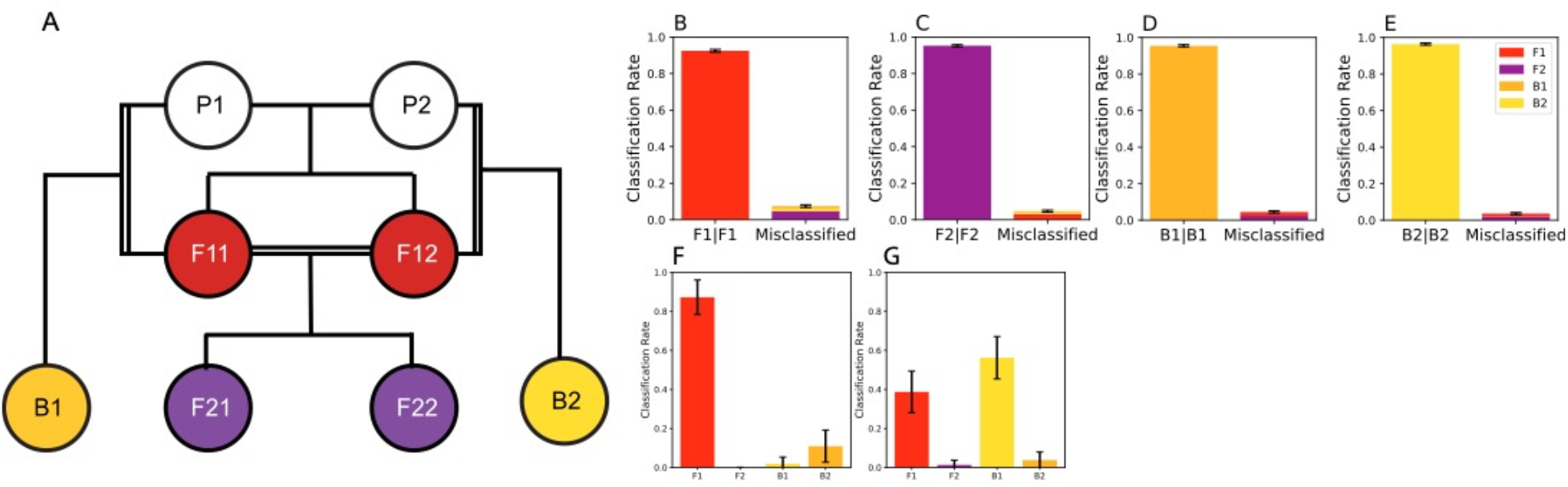
Classification of Backcrossed, F1, and F2 parasites in a typical lab-cross experiment with two rounds of transmission. (**A**) Pedigree used to represent the genealogical history of a laboratory-based genetic cross with two, unrelated parents (P1, P2). F1 progeny are FS and represented by nodes F11 and F12, F2 progeny by nodes F21 and F22. Backcrossed progeny to P1 are represented by B1 and backcrossed progeny to P2 are represented by B2. Inbreeding is denoted by double lines. True classification and misclassification rates for (**B**) F1, (**C**) F2, (**D**) B1, and (**E**) B2. Inferred classifications in the two-generation cross for the **F**) first and **G**) second generation progeny pools. The error bars represent two standard errors from the mean from 40000 simulated comparisons.

Classification involved a two-step triangulation strategy (**Supplemental Text**) that 1) first identified backcrossed progeny and then 2) disambiguated F1 and F2 progeny. Importantly, because the pedigree was known, the likelihood used in *MalKinID* could be modified (**Equation S2-S3**) to infer genealogical relationships based on three-way comparisons (trios) to triangulate the exact nodal position of a sample in the pedigree tree (**Figure 6A, Supplemental Figure 8**) (Sieberts *et al*. 2002). *In silico* experiments show that this approach identified simulated F1, F2, and backcrossed progeny with > 90% accuracy (**Figure 6B-E**). The true classification rate for F1 progeny was 0.925 (95% CI 0.917, 0.932), F2 progeny was 0.953 (95% CI 0.947, 0.959), B1 progeny 0.954 (95% CI 0.949, 0.960), and B2 progeny 0.964 (95% CI 0.958, 0.969).

When applied to the empirical data, it correctly identified 0.87 (95% CI 0.78, 0.96) of the non-parental strains in the first-generation pool as F1’s (**Figure 6F**). The data from this two-generation cross were not used to calibrate the meiosis model and was independent from the previously mentioned NF54 x NHP4026 cross used to calibrate the meiosis model. When applied to the second-generation pool, *MalKinID* showed that 0.39 (95% CI 0.28, 0.49) of the second-generation progeny were F1s, 0.56 (95% CI 0.45, 0.67) were backcrosses involving NF54, and that only a very small fraction, 0.0125 (95% CI 0, 037) were F2s (**Figure 6G**).

## Discussion

*MalKinID* is a likelihood-based model that infers the genealogical relationships between parasites based on the total, genome-wide proportion of IBD shared between individuals and two aspects of the IBD segment block distribution. *MalKinID* accurately identifies first-, second-, and third-degree relatives from one another and can further distinguish the sub-relationships within first-degree relatives (PC, FS, and MS). Importantly, these relationships can be identified even in the presence of meiotic siblings (**Supplemental Figure 6**), despite the fact their expected relatedness (0.33) lies between the expected relatedness of the other first-degree relatives (0.50) and second-degree relatives (0.25). Genealogical inference with *MalKinID* is most powered when: 1) when outcrossing predominates, or 2) when a pedigree specifying the genealogical history connecting individuals can be specified. When these conditions are satisfied, *MalKinID* can accurately reconstruct parasite genealogical histories to evaluate potential transmission histories.

These constraints could pose a challenge when evaluating parasites in natural populations because the genealogical history connecting related parasites is unknown and because inbreeding can be common in low transmission regions (Neafsey and Volkman 2017; Neafsey *et al*. 2021). Inbreeding is a known problem with genealogical inference (Kirkpatrick *et al*. 2011) but is especially problematic for malaria. Many of the genealogical inference techniques used to account for inbreeding were designed for diploid organisms (Liu *et al*. 2010) and do not work in malaria because its predominantly haploid genome prevents using the IBD between homologous chromosomes to identify inbred individuals (Jacquard 1975).

To limit the effects of inbreeding, one can use *MalKinID* to study genetically related parasites from different populations. Parasites sampled from different populations are less likely to be inbred and inter-population genetic relatedness analyses have been used to generate different hypotheses regarding importation (Taylor *et al*. 2017). However, these studies typically focus on the most highly related parasite pairs (*r*_*total*_ > 0.5) (Taylor *et al*. 2017) and do not utilize any additional information from the IBD segment block distribution. Genealogical inference with *MalKinID* would complement previous studies by enabling more granular investigations based on identifying PC, GC, and GGC relatives to identify imported parasites, identify their most likely source population, and estimate the number of rounds of local transmission following importation.

An alternative solution would be to avoid relying on pairwise comparisons and instead focus on utilizing multi-sample comparisons based on a hypothesized pedigree tree that describes the genealogical history of genetically related parasites. This approach overcomes inbreeding limitations by leveraging the relatedness among multiple samples to assess the joint likelihood across all examined sample-pair comparisons, effectively shifting the focus of genealogical inference from identifying the most likely pairwise relationship to the most likely pedigree tree that best explains the relatedness within a sample cluster. Tree-based genealogical inference helps constrains genealogical inference and could improve robustness against minor errors in identifying IBD segments. Future iterations of *MalKinID* will focus on combining the likelihood function with a random pedigree tree simulator (Ochoa; Huisman 2017) to enable efficient tree- based likelihood maximation and genealogical inference.

Multi-sample and tree-based genealogical inference was especially useful for dissecting the genealogical relationships of laboratory-based genetic cross progeny (Vendrely et al. 2020; Button-Simons et al. 2021b; Kumar et al. 2022b). Interestingly, *MalKinID* showed the majority of second-generation progeny were actually F1s or backcrosses to the NF54 parental strain, which could indicate preferential mating with NF54 or the result of high rates of NF54 selfing in the first generation. Using *MalKinID,* to identify and classify recombinant, inbred parasites could be used to better inform genetic linkage mapping and QTL analyses (Daley and Shepherd 2008; Solberg Woods 2014), disentangle the contributions of individual mutations and genetic background to clinically relevant phenotypes such as drug resistance, and study other traits related to virulence, invasion, or transmission (Vendrely *et al*. 2020b).

A critical aspect of *MalKinID* is that it treats each of the 14 per-chromosome IBD segment block distributions as statistically independent. This is important, because selection can distort the IBD segment block distribution and cause longer than expected IBD segment blocks to appear.

However, the effects of selection tend to be localized (Charlesworth *et al*. 2000; Schaffner *et al*. 2018) and are unlikely to extend to other chromosomes due to independent reassortment. As a result, *MalKinID* should accurately distinguish genealogical relationships even if certain regions of the genome are undergoing strong, directional selection. This assumption is partially validated by the high accuracy of *MalKinID* on the empirical lab-cross data, which previously showed evidence of selective sweeps due to culture-adaptation (Vendrely *et al*. 2020a; b). One way of correcting for any potential distortions would be to remove chromosomes with evidence of selective sweeps from analyses (Guo *et al*. 2024).

When running *MalKinID,* it may be advantageous to exclude meiotic siblings (and their descendants, FAV.MS and FCS.MS) as their inclusion could unnecessarily depress true classification rates in situations where meiotic siblings are expected to be rare. Ideally, future studies should incorporate a Bayesian prior to *MalKinID* that specifies the probability that a meiotic sibling will be present in the examined dataset. In the absence of data to inform this prior, the following heuristic is advised. When evaluating parasites in situations where it is highly unlikely that parasites will be derived from the same oocyst, such as when examining parasites between monogenomic (single strain) infections that are the progeny from lab cross experiments (**Figure 4G**), meiotic siblings and their derivatives can be excluded. When examining parasites within polygenomic infections, particularly those that are the result of co-transmission, meiotic siblings and their derivatives should not be excluded when making inferences with *MalKinID* as there is a greater chance of comparing parasites derived from the same oocyst. Note that polygenomic infections will need to first be deconvoluted using technologies such as long-read sequencing (Sakamoto *et al*. 2022) or single-cell (Nair *et al*. 2014; Nkhoma *et al*. 2020) whole genome sequencing to establish the genomic phase of each co-infecting strain phase prior to analysis.

In conclusion, *MalKinID* is a new likelihood-based classification model that can be used to identify the genealogical relationships of genetically related parasites. In theory, *MalKinID* can be used to identify the parent-offspring relationships in natural parasite populations. Pairwise relatedness-based inferences of genealogical history work best in for identifying importations in predominantly outbreeding populations where inbreeding is infrequent. When inbreeding is present, relatedness-based genealogical inference will need to focus on utilizing multi-sample, tree-based inferences to accurately identify the genealogical relationships between parasites.

## Methods

### Meiosis simulation

Genetically related progeny were simulated using a previously published meiosis model that incorporates obligate chiasma formation and crossover interference (Wong *et al*. 2018). Briefly, the model relies on two parameters, the crossover rate (*k*) that quantifies the kilobase pairs per centimorgan and a crossover interference parameter (*v*) that determines how closely crossover events can be placed next to one another on a four-chromatid bundle. A full specification of the meiosis model can be found in our previously published paper (Wong *et al*. 2018) and a brief description of the meiosis model, along with full details regarding the meiosis model recalibration are presented in **Supplemental File 1**.

### Likelihood model structure

A likelihood model was used to determine the most likely genealogical relationship between parasite pairs based on 1) the total genetic relatedness (*r*_*total*_), 2) the size of the largest IBD segment per chromosome (*IBD*_*max*_), and 3) the number of IBD segments per chromosome (*n*_*segments*_). Both *r*_*total*_ and *IBD*_*max*_ are bound between 0 and 1 because *r*_*total*_ was defined as the fraction of the genome that is IBD and *IBD*_*max*_ was normalized and defined as the proportion of the chromosome the segment sits on. The structure of the likelihood model (**Equation 1**) was defined as:

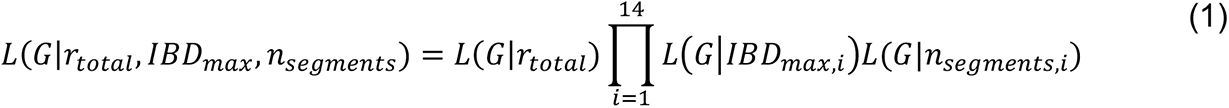

where *i* refers to the *i-*th chromosome in the genome and *G* refers to the examined genealogical relationship (**Table 1**).

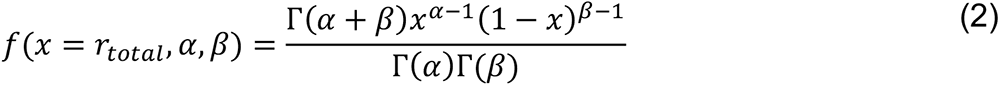

whose parameters, α and β, were fit to the simulated *r*_*total*_ values using the *scipy.stats.beta.fit* function from the scipy package (v 1.7.3) for Python 3. The location and scale parameters for the *scipy.stats.beta.fit* function were set to 0 and 1, respectively.

(𝐺|*IBD*_*max*,i_) was defined using a Kernel density estimator (KDE) with spikes (Tataru *et al*. 2015; Guerrero Montero and Blythe 2023) and defined as:

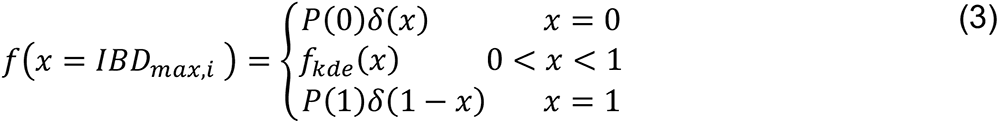

where P(0) is the probability that *IBD*_*max*,i_ is zero (no IBD segment on chromosome *i*), P(1) is the probability that *IBD*_*max*,i_ is 1 (the IBD segment spans the length of chromosome *i*), 𝛿 is a Dirac delta function, and 𝑓_𝑘𝑑𝑒_ is a KDE that was fit to the remaining data where *IBD*_*max*,i_ > 0 and *IBD*_*max*,i_ < 1 using the *KernalDensity* function in the Python 3 package *sklearn.neighbors* (v1.2.2).

(𝐺|*n*_*segments*_) was defined as probability mass function (PMF) with add-one additive smoothing:

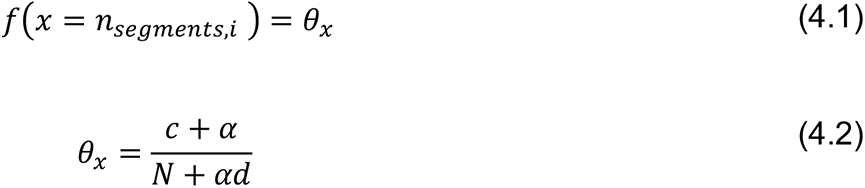

where *θ*_*x*_ is the smoothed probability of observing *n*_*segments,i*_ and 𝛼 is a pseudocount used for smoothing and set to 1. N, the number of trials examined, *d,* the number of *n*_*segment*𝑠,i_ categories, and *c*, the number of simulations where *n*_*segments,i*_=x. N was set to 5000 (the number of simulations run) and *d* was the number of simulated categories for *n*_*segments,i*_ plus 1. The last index of *d* represents a miscellaneous category for any segment count that was not observed in the simulation.

### Classification

Classification was based on maximum likelihood analysis. The classification with the highest likelihood was chosen as the most likely hypothesis. For composite hypotheses, the general likelihood ratio test was used:

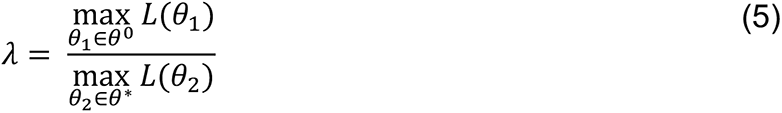

where *θ*_1_ is a null hypothesis belonging to a set of null hypotheses in *θ*^8^ and *θ*_7_ is an alternative hypothesis belonging to a set of alternative hypotheses in *θ*^∗^.

### Laboratory-cross data

Whole genome sequences for the progeny from three laboratory-cross (MKK2835 x NHP1337, NF54 x NHP4026, and Mal31 x KH004) as described in (Vendrely et al. 2020; Button-Simons et al. 2021b; Kumar et al. 2022b). The whole genomes sequences for each set of progeny were filtered to include only sites that were variant between the two parental lines, were biallelic, had an allele depth >10, and had less than 20% of the sites missing across all the samples collected from that cross. This left 7177 SNP variants for the MKK2835 x NHP1337 cross, 11358 variants for the NF54 x NHP4026 cross, and 12918 variants for Mal31 x KH004 cross.

For the cross data, IBD was inferred using a modified version of the *hmmIBD* (Schaffner *et al*. 2018) as described previously in (Wong *et al*. 2018). This modified version replaces the emission probabilities with:

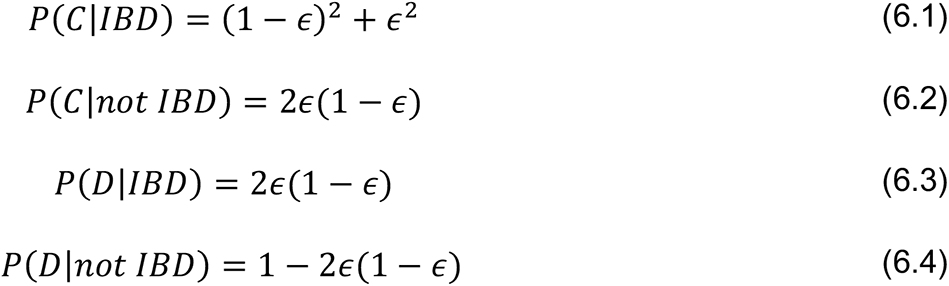

where C indicates concordance, D indicates discordance, and 𝜖 is the sequencing error rate and set to 0.01. After running the modified *hmmIBD*, IBD segments were identified as contiguous blocks of loci inferred to be IBD. Segments whose length was less than 0.05 of the chromosome on which it resided were removed from analysis. These spurious IBD segments could represent spurious artifacts generated by *hmmIBD* or mitotic recombination events. Total relatedness was quantified as the total proportion of the genome that is IBD.

### F1-score

The F1-score is the harmonic mean of the precision (the positive-predictive-value) and sensitivity (recall). It was defined as:

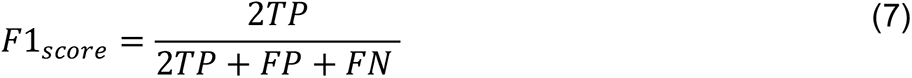

where TP is the number of true positives, FP the number of false positives, and FN the number of false negatives.

## Acknowledgements

Experimental genetic crosses used were funded by the National Institutes of Health (5P01AI127338-07). Work in the Texas Biomedical Research Institute was conducted in facilities constructed with support from Research Facilities Improvement Program grant C06 RR013556. Model calibration and *MalKinID* development was supported from the Bill and Melinda Gates Foundation (OPP1156051 to DFW and INV-049909 to SKV) and the National Institutes of Health (5R21AI141843-02 to SKV).

---

**Supplemental Figure 1.**
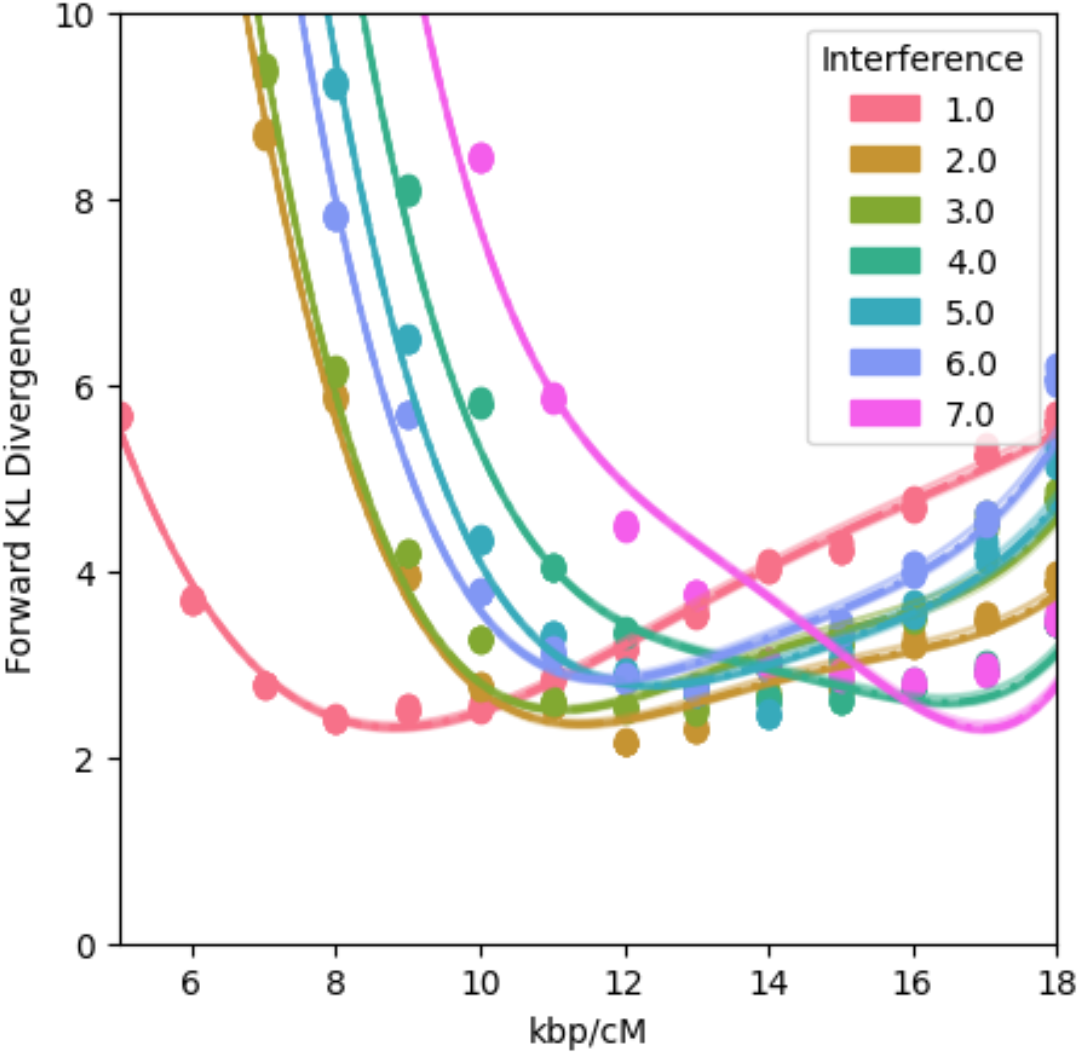
The forward Kullback Liebler divergence (KL divergence) for simulations run with different crossover rates (measured in kilobase pairs per cM) and crossover interference levels. An interference of 1.0 corresponds to no crossover interference. Each dot represents the forward KL divergence obtained after jackknife-by-block resampling the data into 10 equally sized bins. The curves were generated by fitting a four-dimensional polynomial function to each of the jackknife-by-block resampled estimates.

**Supplemental Figure 2A-G.** See Supplemental File 2

**Supplemental Figure 3.**
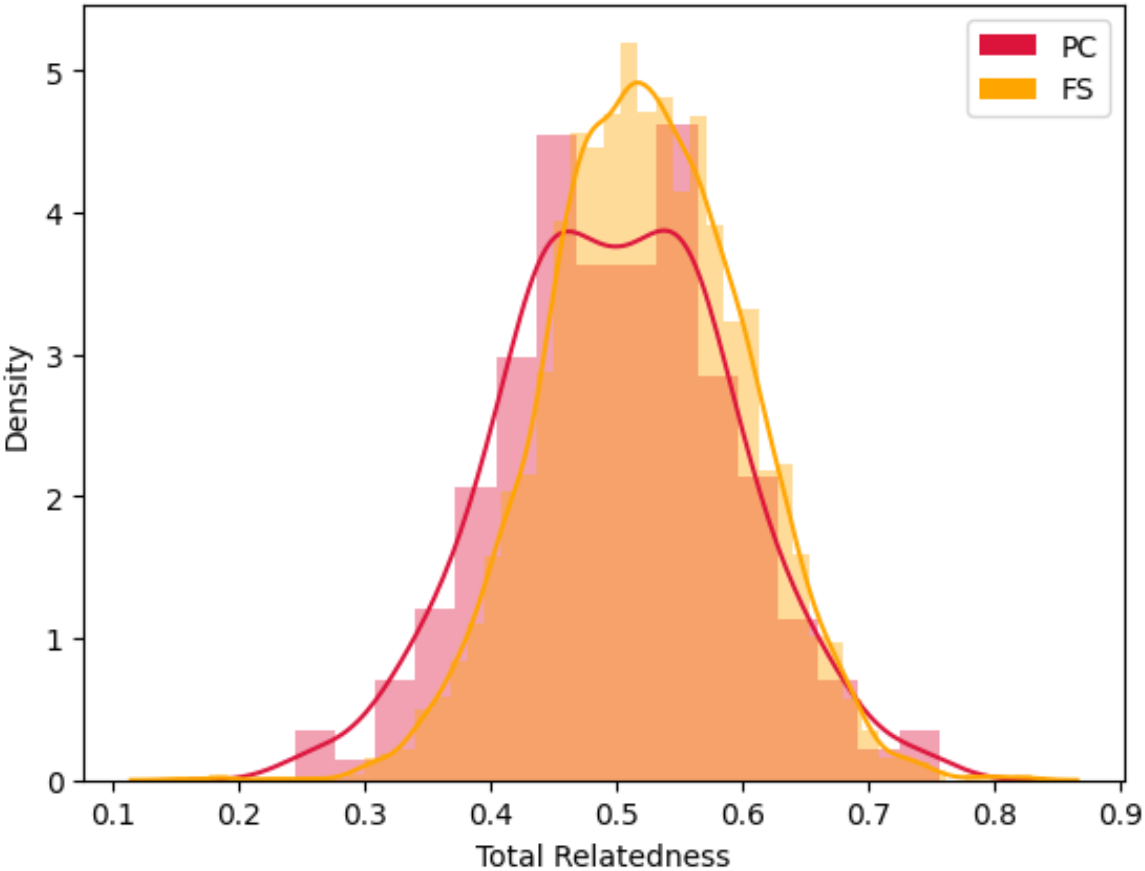
Distribution of pairwise PC and FS relatedness observed in the cross data. The raw average and standard deviation of relatedness for PC was 0.50 and 0.09. The raw average and standard deviation of relatedness for FS was 0.53 and 0.079.

**Supplemental Figure 4.**
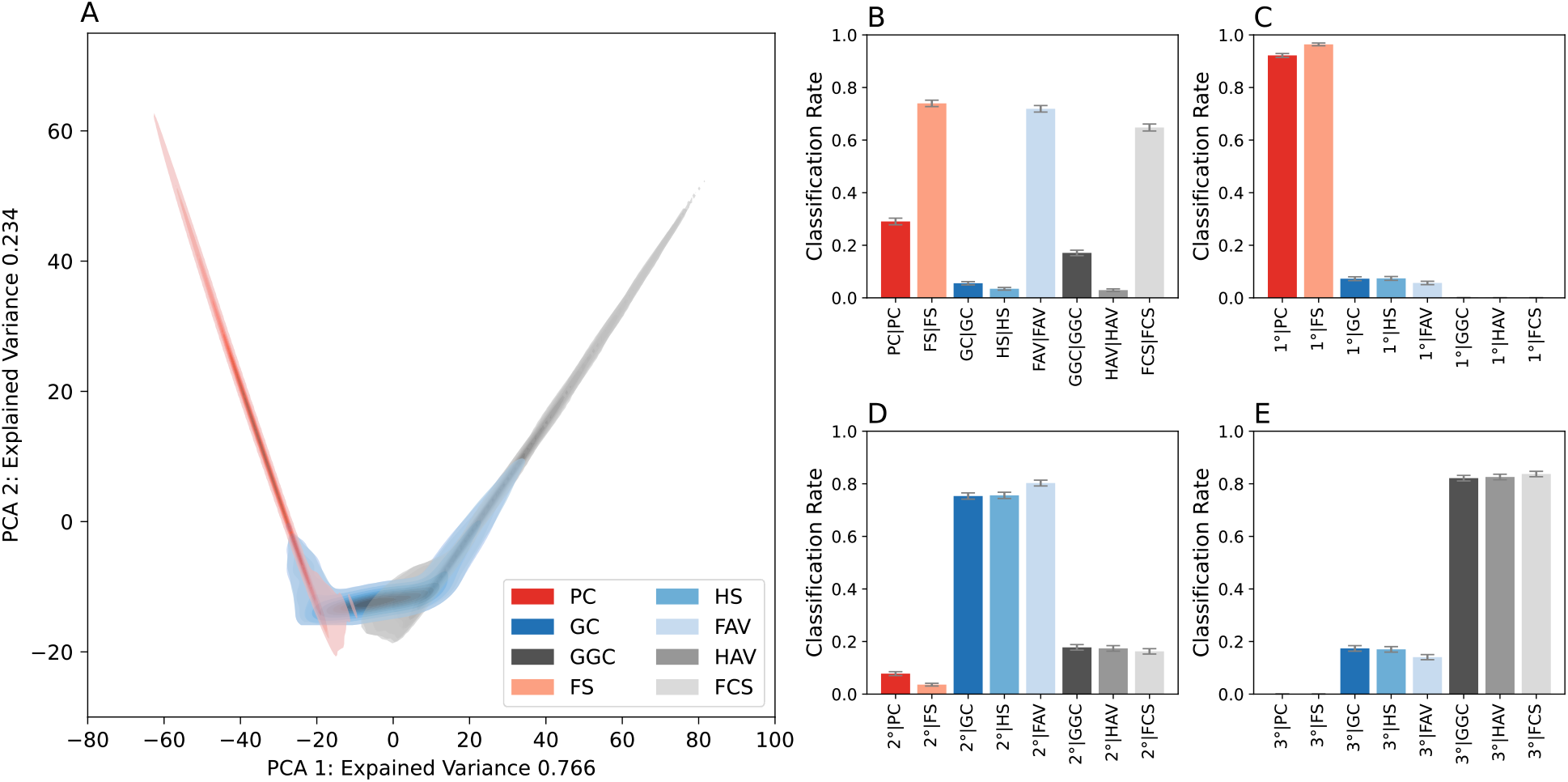
PCA plot of the simulated *r*_*total*_ data. The PCA included a third component that is not shown. Each genealogical relationship is represented by 5000 independent simulations and was fit with a gaussian density estimator for visual clarity. Darker spots indicate a region with higher mass for that genealogical relationship. *MalKinID* classification rates on simulated data using the *r*_*total*_ alone (**B-E**). The notation used in the graphs is similar to that used for conditional probability. The term to the left of the “|” refers to the inferred classification and the term to the right the true genealogical relationship. **B** classifies samples by their genealogical relationship while the other subplots classify samples (**C**) first-, (**D**) second-, or (**E**) third-degree relatives.

**Supplemental Figure 5.**
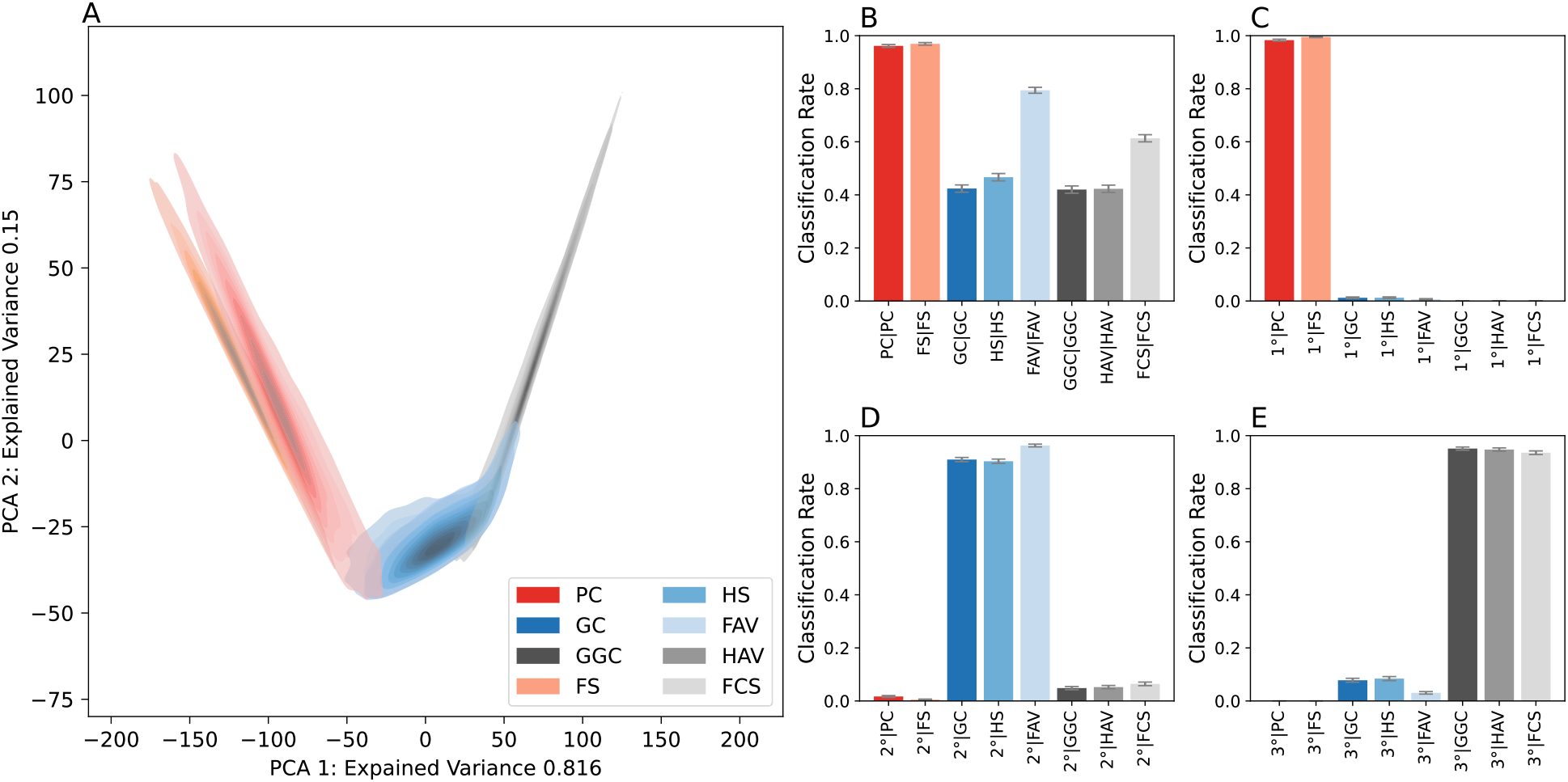
**A**) PCA plot of the simulated *r*_*total*_, *IBD*_*max*_, and *n*_*segment*_data. The PCA included a third component that is not shown. Each genealogical relationship is represented by 5000 independent simulations and was fit with a gaussian kernel density estimator for visual clarity. Darker spots indicate a region with higher mass for that genealogical relationship. *MalKinID* classification rates on simulated data using the joint-likelihood that includes *r*_*total*_, *IBD*_*max*_, and *n*_*segment*_ (**B-E**). The notation used in the graphs is similar to that used for conditional probability. The term to the left of the “|” refers to the inferred classification and the term to the right the true genealogical relationship. **B** classifies samples by their genealogical relationship while the other subplots classify samples (**C**) first-, (**D**) second-, or (**E**) third-degree relatives.

**Supplemental Figure 6.**
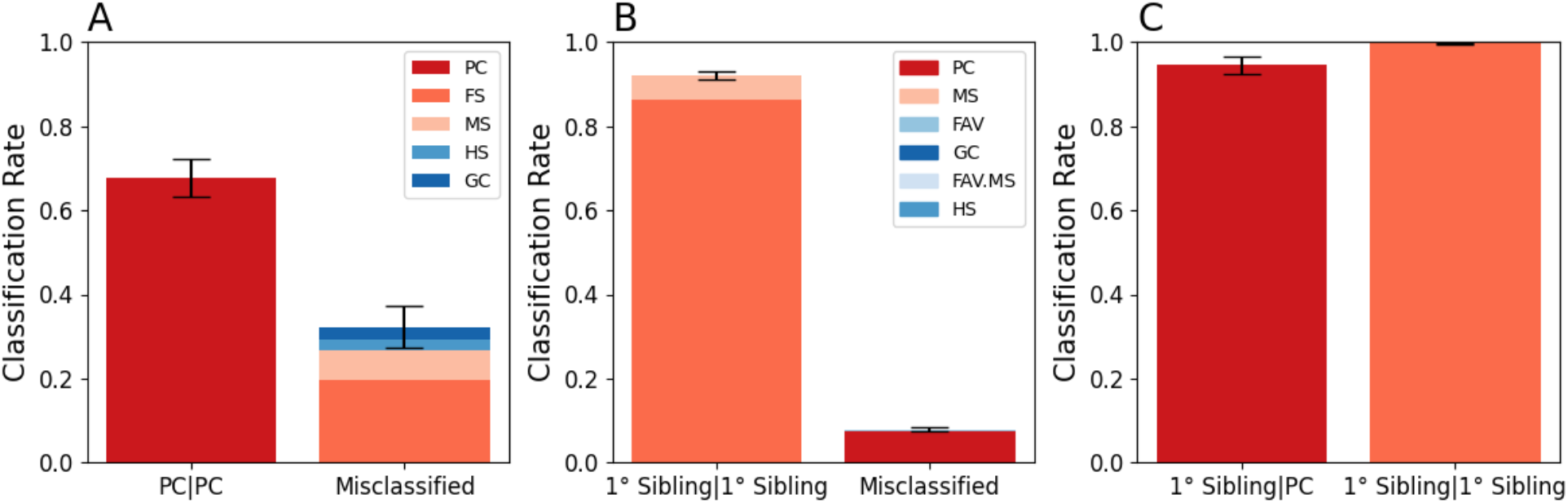
Empirical classification rates in the laboratory cross data using the moderate/high crossover inference parameter set (*v*=7, *k* = 17.3). Error bars indicate two standard deviations from the mean.

**Supplemental Figure 7.**
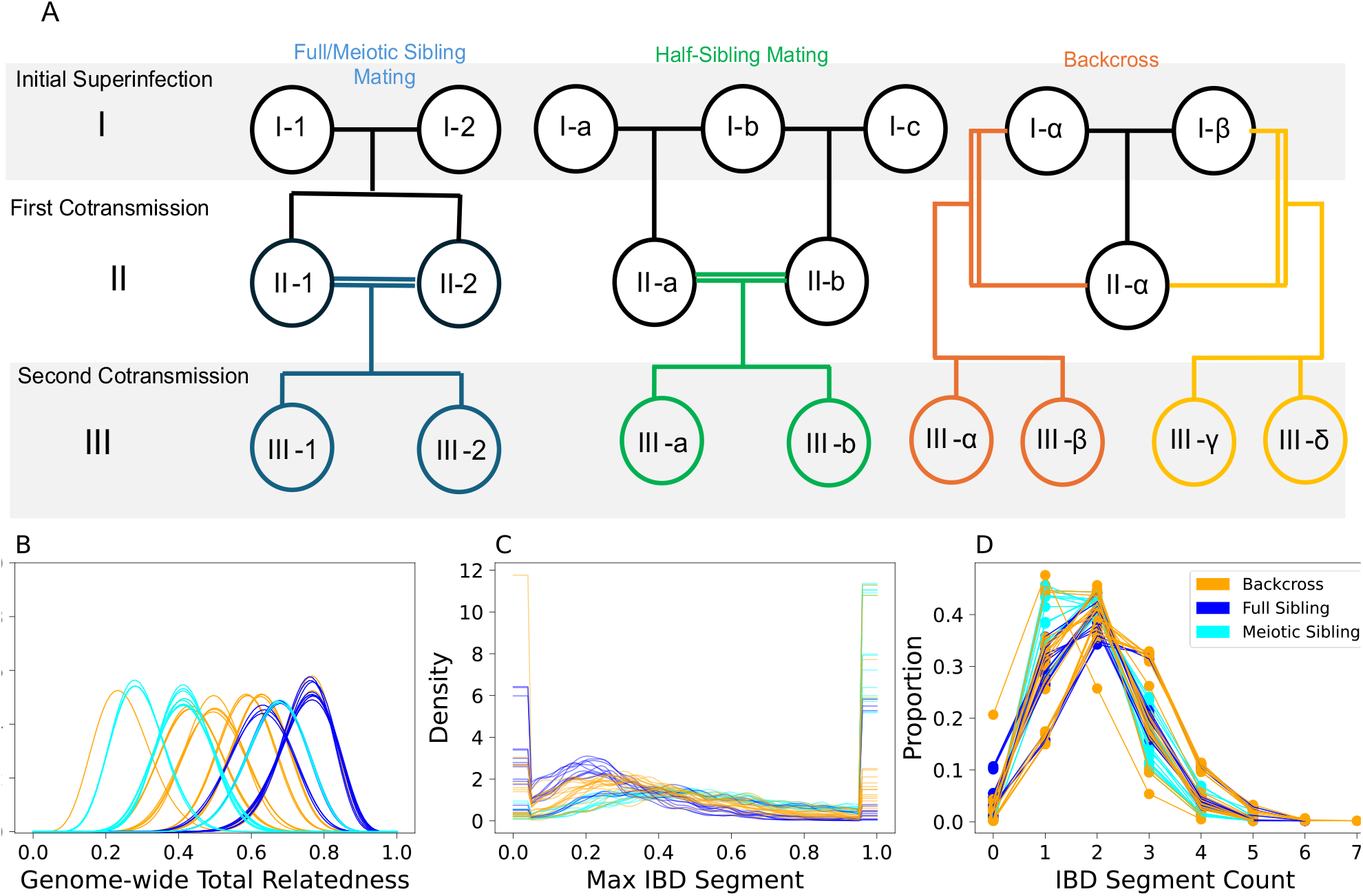
A. The three pedigrees used to represent the types of inbreeding that occur within the first two co-transmission events. The first row specifies the initial, unrelated parental parasite strains that are present in the initial superinfection. The second row are the outcrossed progeny generated after the first co-transmission event, and the third row the inbred progeny of the second co-transmission event. Inbreeding is denoted by double lines. Distributions of (**B**) total genetic relatedness, (**C**) max IBD segment size on chromosome 14, and (**D**) IBD segment count on chromosome 14 for all unique, pairwise comparisons across all three trees involving full siblings (dark blue), meiotic siblings (light blue), and a backcrossed strain (orange).

**Supplemental Figure 8.**
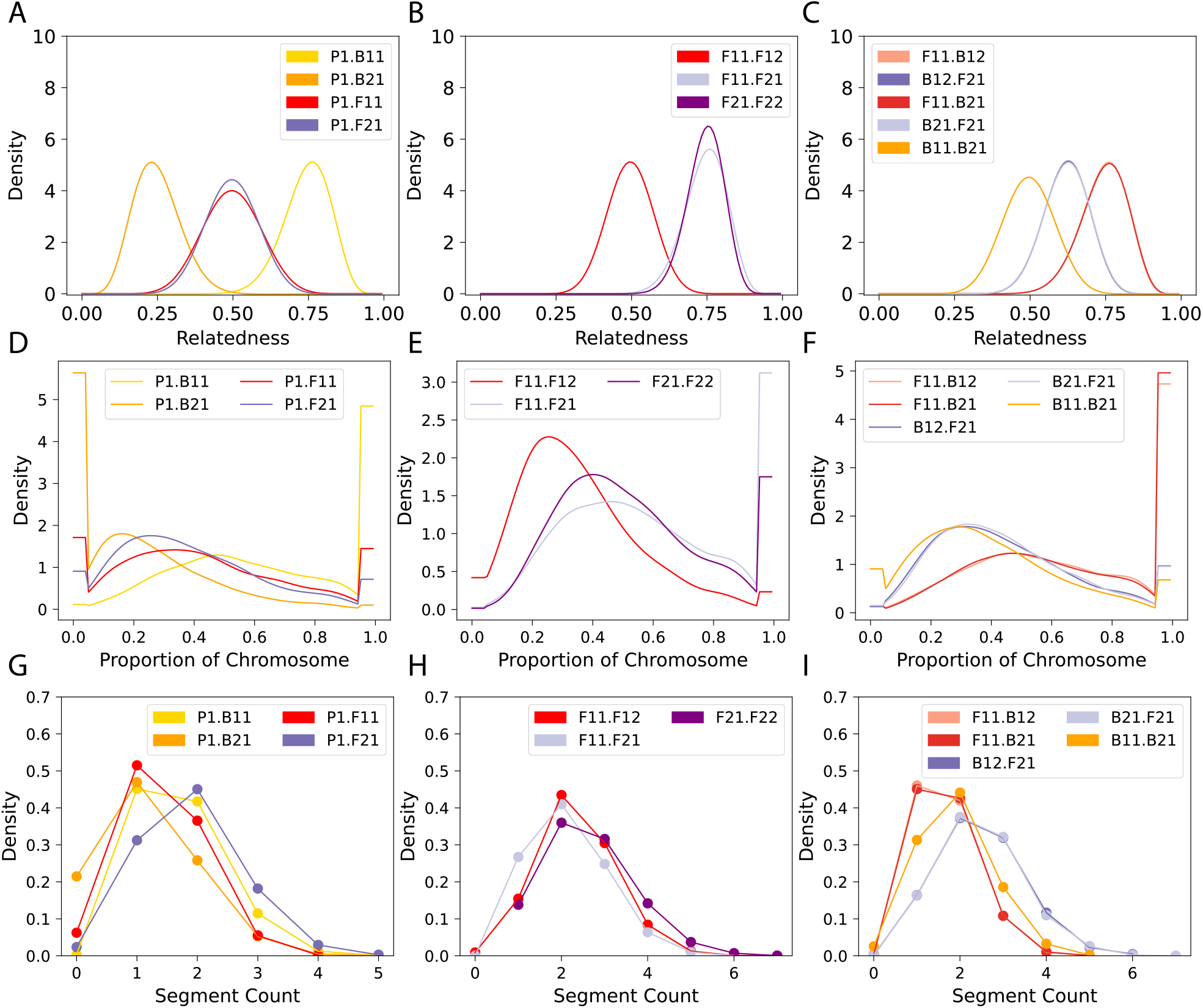
Patterns of genetic relatedness for the laboratory-based crosses. The first column involves comparisons against the first parental strain (P1), the second column involves comparisons between F1 (F11 and F12) and F2 (F21 and F22) progeny, and the third column involves comparisons of F1 and F2 progeny with each of the backcrossed strains (B1 [B11, B12] and B2 [B21, B22]). The first row shows the distributions of *r*_*total*_, the second row the distributions of *IBD*_*max*_ on chromosome 14, and the third row the distributions of *n*_*segment*_ on chromosome 14.

**Supplemental Table 1.** A description of the lab crosses used to calibrate the meiosis model

| Cross | Parent 1 | Parent 1 Origin | Parent 2 | Parent 2 Origin | Number of Unique Progeny | Number of PC comparisons | Number of FS comparisons |
| --- | --- | --- | --- | --- | --- | --- | --- |
| NF54 x NHP4026 | NF54 | Lab-culture | NHP4026 | Southeast Asia | 91 | 182 | 4094 |
| MKK2835 x NHP1337 | MKK2835 | Southeast Asia | NHP1337 | Southeast Asia | 35 | 70 | 595 |
| Mal31 x KH004 | Mal31 | Africa | KH004 | Southeast Asia | 94 | 188 | 4371 |

## Data Availability

Whole genome sequencing data for cloned progeny from lab generated *Plasmodium falciparum* crosses used for calibration (NF54 x NHP4026, MKK2835 x NHP1337, and Mal31 x KH004) were previously published and available at NCBI under project PRJNA524855. Filtered SNP genotypes for these crosses and the two-generation crosses involving NF54 x NHP4026 are available at https://github.com/emilyli0325/P01-cloned.progeny.

**Code Availability**: Code to run *MalKinID* will be available on GitHub (https://github.com/weswong/MalKinID) and archived at Zenodo at time of publication.

**Supplemental File 1:** Supplemental Text

**Supplemental File 2:** Supplemental Figure 2 containing all the IBD segment count and max IBD segment distributions for first-, second-, and third-degree relatives.

**Supplemental File 3:** Parameter Tables

