## Supplemental File 1 for "MalKinID: A Likelihood-Based Model for Identifying Malaria Parasite Genealogical Relationships Using Identity-by-Descent"

*Meiosis model description*

Genetically related progeny were simulated using a previously published meiosis model that incorporates obligate chiasma formation and crossover interference [1]. Briefly, each genome is represented by a vector with 23 million base pairs where each element of the vector is defined by an integer that represents the ancestry of that genomic position. For example, an integer value of 0 indicates that position was inherited from parent “0”. This vector is then subdivided into chromosomes based on the chromosome lengths reported in *PlasmoDB*. The model independently simulates meiotic recombination for each of the 14 chromosomes in the *P. falciparum* genome by:

1) Placing an initial chiasma on a four-chromatid bundle by drawing from a uniform distribution that spans the length of the examined chromosome
2) Placing subsequent chiasma upstream and downstream of the initial chiasma by drawing from a constrained gamma distribution where the parameter, $v$, determines the strength of crossover interference until the ends of the chromosome are reached.
3) For each chiasma, a sister chromatid from each parental homolog is randomly selected to undergo recombination
4) After all chiasma are resolved, the model independently segregates and randomly combines sister chromatids from each of the bivalents for each chromosome to create haploid parasite genomes.

IBD was determined by comparing the genomes of each simulated parasite. If the integer values at a given position was identical, it was considered IBD. If not, it was considered not IBD. The meiosis model is specified by two parameters: 1) a crossover interference parameter ($v$) and 2) a crossover rate parameter ($k$, measured in kilobasepairs per centimorgan or kbp/cM).

Together, these two parameters determine the spacing of chiasmata on the four-chromatid bundle. The crossover interference parameter that prevents chiasma from being spaced too closely together. v=1 corresponds to no crossover interference and v>1 corresponds to increasingly high levels of crossover interference.

*Meiosis model recalibration*

The meiosis model was originally calibrated to the parent-child (PC) relationships observed in 67 *P. falciparum* progeny generated from lab-crosses involving (3D7, HB3, Dd2, 7G8, and GB4) [1]. These data were sufficient to show that a model with obligate chiasma formation fit the data better than one without but was insufficient to identify the exact level of crossover interference. For this study, we recalibrated the model to data from three recently performed lab crosses: NF54 x NHP4026 (a sustained, lab-cultured strain crossed with a recently isolated Southeast Asian strain), MKK2835 x NHP1337 (two recently isolated Southeast Asian strains), and Mal31 x KH004 (a recently isolated African crossed with a recently isolated Southeast Asian strain) (**Supplemental Table 1**).

The model was recalibrated to the total relatedness, max IBD segment size per chromosome, and segment count per chromosome from all usable parent-child (PC) and full-sibling (FS) comparisons. Parent-child comparisons were identified as any comparison involving one of the parental strains and all comparisons involving only progeny strain were assumed to be full-siblings. The possibility of meiotic siblings was ignored because the experimental design of the laboratory crosses made it highly unlikely for progeny to have originated from the same oocyst.

Model fit was evaluated using the forward Kullback-Leibler divergence (KL divergence) using the Python 3 *scipy.special.kl_div* function (v1.7.3) and defined as:

$$D_{KL}(P||Q) = \sum_{x \epsilon X} P(x) \log\left(\frac{P(x)}{Q(x)}\right) \tag{S1.1}$$

$$D_{KL}(P||Q_{v,k}) = \sum_{c=1}^{3} D_{KL,c}(P_c||Q_{v,k}) \tag{S1.2}$$

$$D_{KL,c}(P_c||Q_{v,k}) \tag{S1.3}$$

$$= \sum_{G \epsilon \{PS,FS\}} \Big[ D_{KL,c,G}(P_{r_{total,c,G}}||Q_{v,k_{r_{total,G}}}) + \sum_{i=1}^{14} \Big[ D_{KL,c,G}(P_{IBD_{max,c,G,i}}||Q_{v,k_{IBD_{max,G,i}}}) + D_{KL,c,G}(P_{n_{segment,c,G,i}}|| Q_{v,k_{n_{segment,G,i}}}) \Big] \Big]$$

**Equation S1.1** describes the general equation for the forward KL divergence. P is the "true" probability distribution that is going to be approximated by the probability distribution Q. **Equations S1.2** and **S1.3** describe the objective function used for model calibration. In **Equation S1.2**, $P_c$ is the empirical probability distribution obtained from the data in cross *c* and $Q_{v,k}$ is the simulated probability distribution obtained from running the meiosis model 5000 times and setting the crossover interference parameter to *v* and the crossover rate to *k*. *c* is the index of the *c-th* genetic cross in the set {NF54xNHP4026, MKK2835xNHP1337, Mal31xKH004}. Note that this specification assumes that the empirical probability distributions obtained from the cross data are the "true" distribution that need to be approximated by $Q_{v,k}$.

In **Equation S1.3**, $P_{r_{total,c,G}}$ is the empirical probability density function (PDF) for total relatedness for genealogy *G* (which is limited to either PC or FS) using data from cross *c*.

$Q_{v,k}{}_{r_{total,G}}$ is the simulated PDF for total relatedness for genealogy *G* using the meiosis model with parameters *v* and *k*. $P_{IBD_{max,c,G,i}}$ and $P_{n_{segment,c,G,i}}$ are the empirical PDFs for $IBD_{max}$ and PMFs for $n_{segment}$ on chromosome *i* for genealogy *G* using the data from cross *c*. $Q_{v,k}{}_{IBD_{max,G,i}}$ and $Q_{v,k}{}_{IBD_{max,G,i}}$ are the simulated PDFs for $IBD_{max}$ and PMFs for $n_{segment}$ on chromosome *i* for genealogy *G* when running the meiosis model with parameters *v* and *k.* The PDFs associated with $r_{total}$ and $IBD_{max}$ were approximated by setting 10 equally sized bins covering the range from 0 to 1 and calculating the proportion of data falling in each bin. The PMFs associated with $n_{segment}$ were approximated by setting 10 bins, where the index of each bin is used to define the IBD segment count category, and calculating the proportion of data falling in each bin.

For the empirical data, IBD was inferred using a modified version of the *hmmIBD* [2] as described previously in *Wong et al* [1]. This modified version replaces the emission probabilities with:

$$P(C|IBD) = (1-\epsilon)^2 + \epsilon^2 \quad \text{(S2.1)}$$

$$P(C|not\ IBD) = 2\epsilon(1-\epsilon) \quad \text{(S2.2)}$$

$$P(D|IBD) = 2\epsilon(1-\epsilon) \quad \text{(S2.3)}$$

$$P(D|not\ IBD) = 1 - 2\epsilon(1-\epsilon) \quad \text{(S2.4)}$$

where C indicates concordance, D indicates discordance, and $\epsilon$ is the sequencing error rate and set to 0.01. Prior to running this modified *hmmIBD,* the data from each cross were filtered to include only SNPs that differ between the two parental strains.

After running the modified *hmmIBD*, IBD segments were identified as contiguous blocks of loci inferred to be IBD. Segments whose length was less than 0.05 of the chromosome it resided on were removed from analysis. These spurious IBD segments could represent spurious artifacts generated by *hmmIBD* or mitotic recombination events. Total relatedness was quantified as the total proportion of the genome that is IBD.

*Choosing the optimal meiosis model parameterization*

The optimal model parameterization for the meiosis model was determined using a two-parameter sweep across different crossover interference levels $v \epsilon \{1..7\}$ and crossover rates $k \epsilon \{5..18\}$ and minimizing the forward KL divergence (**Equation S1.1-S1.3**). Model calibration included jackknife-by-block-resampling that randomly split the data from the PC and FS data from the crosses into 10 equally sized bins, calibrating the model to 9 of them, and repeating the process until each bin has been excluded at least once. For each set of jackknife-by-block resampled estimates, a polynomial curve with four dimensions was fit to the estimated forward KL divergences using the Python 3 numpy.polyfit (v1.21.5) function. The resulting curve was then used to identify the parameter combination that minimized the forward KL divergence.

The parameter sweep identified multiple local minima (**Supplemental Figure 1**): one with no crossover interference (v=1, k = 8), one with weak crossover interference (v=2, k = 11.3), and one with moderate/high crossover interference (v=7, k=17). Of these, the parameter set with no crossover interference (v=1, k=8) was excluded because crossover interference is a fundamental feature of meiosis and has been observed across multiple taxa. For the remaining parameter choices, we evaluated their true PC and FS classification rates by fitting the likelihood model (**Methods, Equation 1-4**) to the simulated data generated using each parameter choice and using the resulting likelihood model to classify the data from the three empirical crosses.

Overall, the total true classification rate (defined as the average of the true PC and FS classification rates) for both parameter sets was 0.82. However, the two parameterizations had different true PC classification and true FS classification rates (**Supplemental Figure 7**). The weak crossover interference parameterization (v=2, k=11.3) had a true PC classification rate of 0.863 and a true FS classification rate of 0.781. The moderate/high crossover interference parameterization (v= 7, k = 17) had a true PC classification rate of 0.73 and a true FS classification rate of 0.90.

For pedigree reconstruction and eventual transmission tree reconstruction, we reasoned that maximizing the true classification of vertical relationships (PC, GC, GGC) was more useful, as these relationships most informative for reconstructing genealogical and transmission history. Based on this logic, the weak crossover interference parameterization (v=2, k=11.3) was chosen for all our simulations.

**Classification of F1, F2, and backcross parasites in lab-cross experiments**

Classification of inbred parasite lines is possible when the structure of the pedigree is known, such as during a laboratory-based cross experiment where the parental strains are known. Because these lab-based crosses are usually optimized to favor mosquito infection and promote oocyst development, we assumed that the resulting progeny are derived from separate oocysts and that it was unlikely for two, genetically distinct parasites to be meiotic siblings. Together, these conditions help constrain genealogical inference and limit classification to four classes: parental strain (P1 or P2), F1 progeny, F2 progeny, and backcrossed progeny (B1 for an F1 backcrossed to P1 and B2 for an F2 backcrossed to P2). *A priori* knowledge of the pedigree also allows us to shift our focus from identifying the pairwise relationship between each strain and instead focus on the nodal placement of each strain within the pedigree.

After the first transmission event, the resulting parasite pool (referred to here as the first progeny pool) consists of a mix of parental strains and F1-progeny. The parental strains in the first progeny pool are the offspring of oocysts that are result from parental selfing (mating with a clonal parasite). The parasites in the second progeny pool (the descendants of the second transmission event) are a mix of parental clones, F1 progeny, F2 progeny, and backcrossed strains. Parental clone identification is trivial and can be achieved by determining whether the genome of the progeny strain matches either of the parents. Identification of F1, F2, and backcross strains is more involved, and we use a two-step classification scheme that first identifies backcrosses and then distinguishes F1 and F2 progeny.

The first part of our classification scheme is to identify backcrossed progeny from F1/F2 progeny based on comparisons to P1 and P2 (**Supplemental Figure 8, first column)**. This takes advantage of the fact that backcrossed progeny are more closely related to the backcrossed parent (*ie* P1 and B1 vs P2 and B2) and that the distributions of $r_{total}$, $IBD_{max}$, and $n_{segment}$ for parental-to-backcross comparisons are different from those for parental-to-F1 or parental-to-F2 comparisons. The joint-likelihood function for this initial, pairwise comparison is defined as:

$$L(G) = \prod_{i=1}^{2} L(G|r_{total,i\to G})L\big(G\big|IBD_{count,i\to G}\big)L(G|IBD_{max,i\to G}) \quad \text{(S3)}$$

where i$\epsilon${P1, P2}, G$\epsilon${B1, B2, F1, F2}, and i->G describes the comparison being examined (*ie* P1-B1, P1-B2, etc). The most-likely classification is then identified using a general likelihood ratio test (**Equation 5**) where the alternative hypothesis is a composite hypothesis of all the other relationships in G that are not currently being examined.

Once the B1 and B2 samples are identified, they are removed and we assume that the remaining samples consist of F1 and F2 progeny (referred to as the F1/F2 pool) that need to be disambiguated. When identifying the classification of a given sample, we examine trios consisting of P1 and two samples (*s1* and *s2*) that are sampled from the F1/F2 pool. By comparing these trio sets, we can identify *s1* based on the differences in $r_{total}$, $IBD_{max}$, and $n_{segment}$ between P1 and F1 progeny, P1 and F2 progeny, two F1 progeny, two F2 progeny, and an F1 and F2 progeny (**Supplemental Figure 8,** column 1 and column 2). The joint-likelihood function is defined as:

$$L(G_{s1:s2}) = \left(G \middle| r_{total,s1:s2},\ IBD_{count,s1:s2}, IBD_{max,s1:s2}\right) * \prod_{i=1}^{2} L\left(G \middle| r_{total,P1:s_i},\ IBD_{count,P1:s_i}, IBD_{max,P1:s_i}\right) \qquad \text{(S4)}$$

where $G \epsilon \{F1\!:F1,\ F1\!:F2,\ F2\!:F1, F2\!:F2\}$, where the first part to the left of the colon is the relationship for *s1* and the part to the right the relationship for *s2*. The joint-likelihood simultaneously infers the genealogical relationship of *s1* and s2.

When applied to the empirical, two-generation cross data, **Equation S4** was applied to all possible combinations of s1 and s2 and the most common G was used as the inferred G for that sample.
