## Supplemental File 2 for "MalKinID: A Likelihood-Based Model for Identifying Malaria Parasite Genealogical Relationships Using Identity-by-Descent"

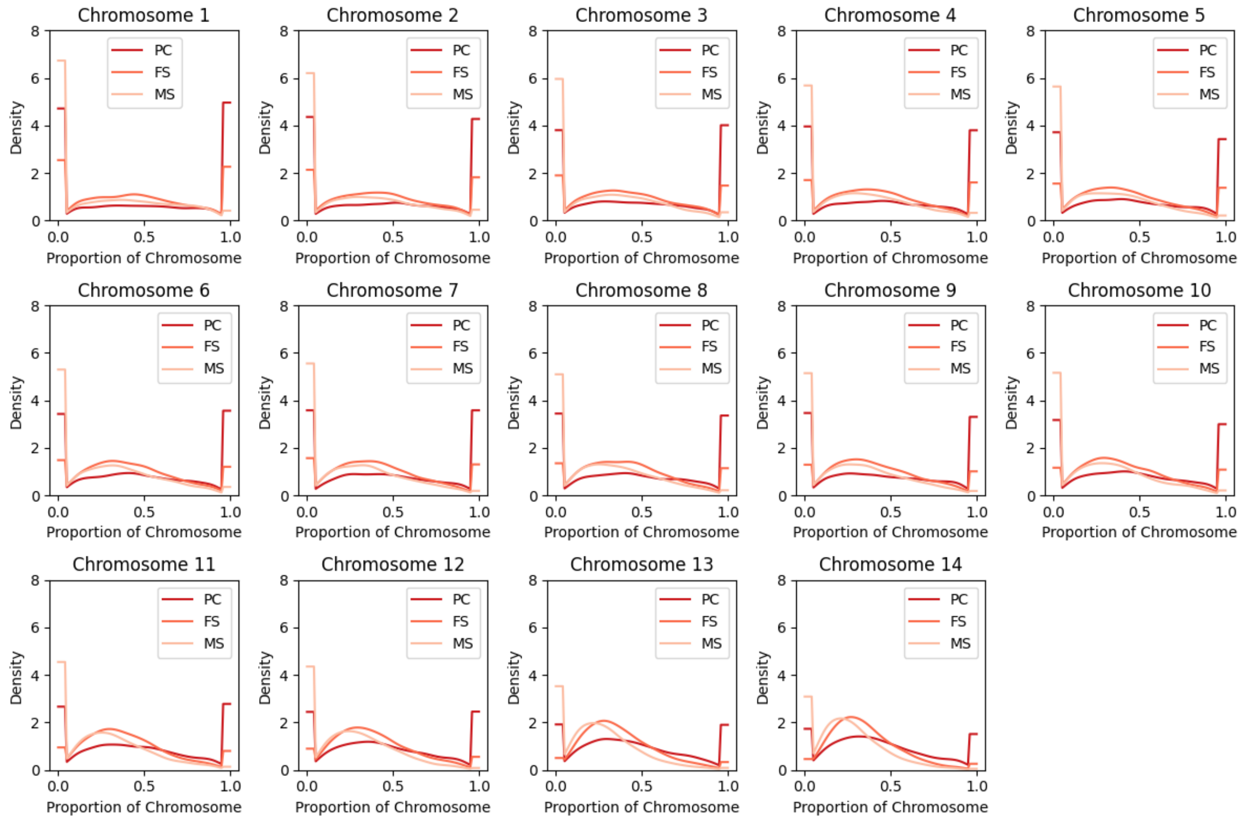


**Supplemental Figure 2A** Simulated max IBD segment block distribution for first-degree relatives.

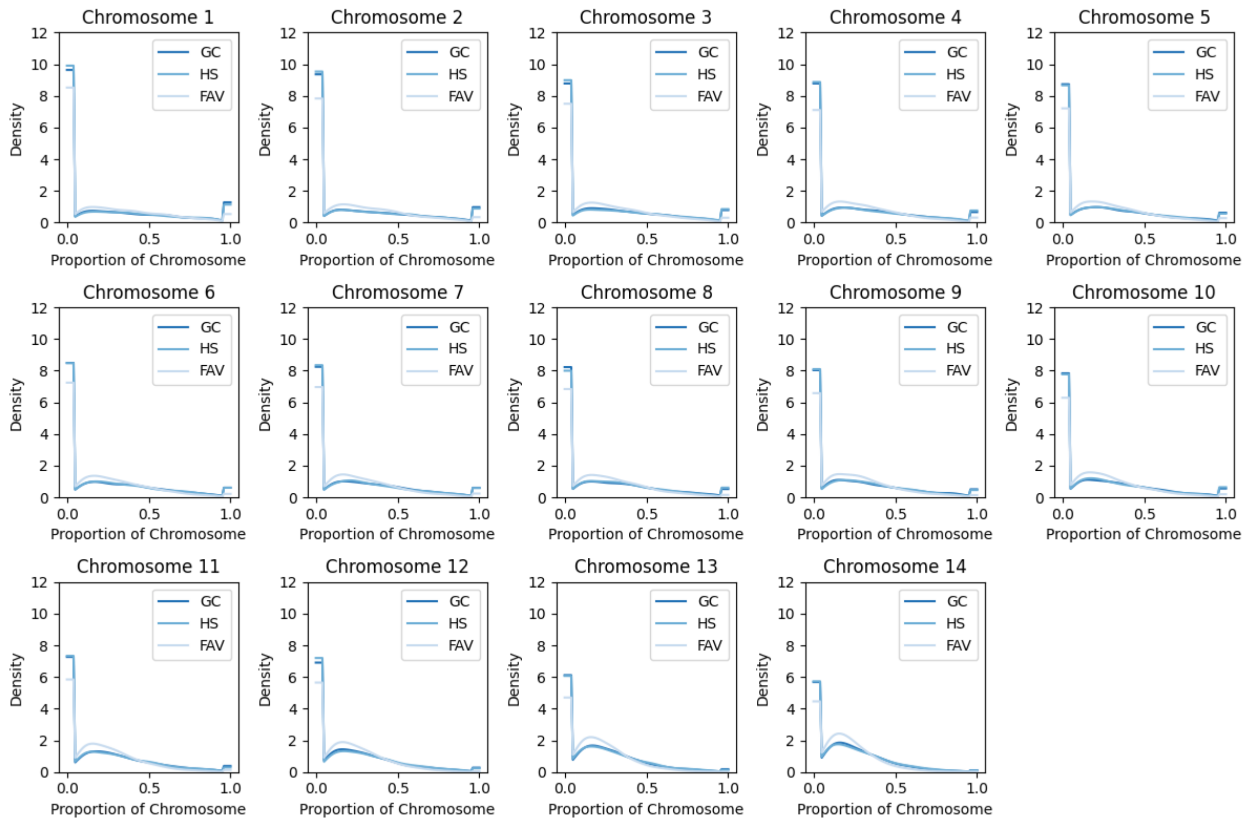


**Supplemental Figure 2B** Simulated max IBD segment block distribution for second-degree relatives.

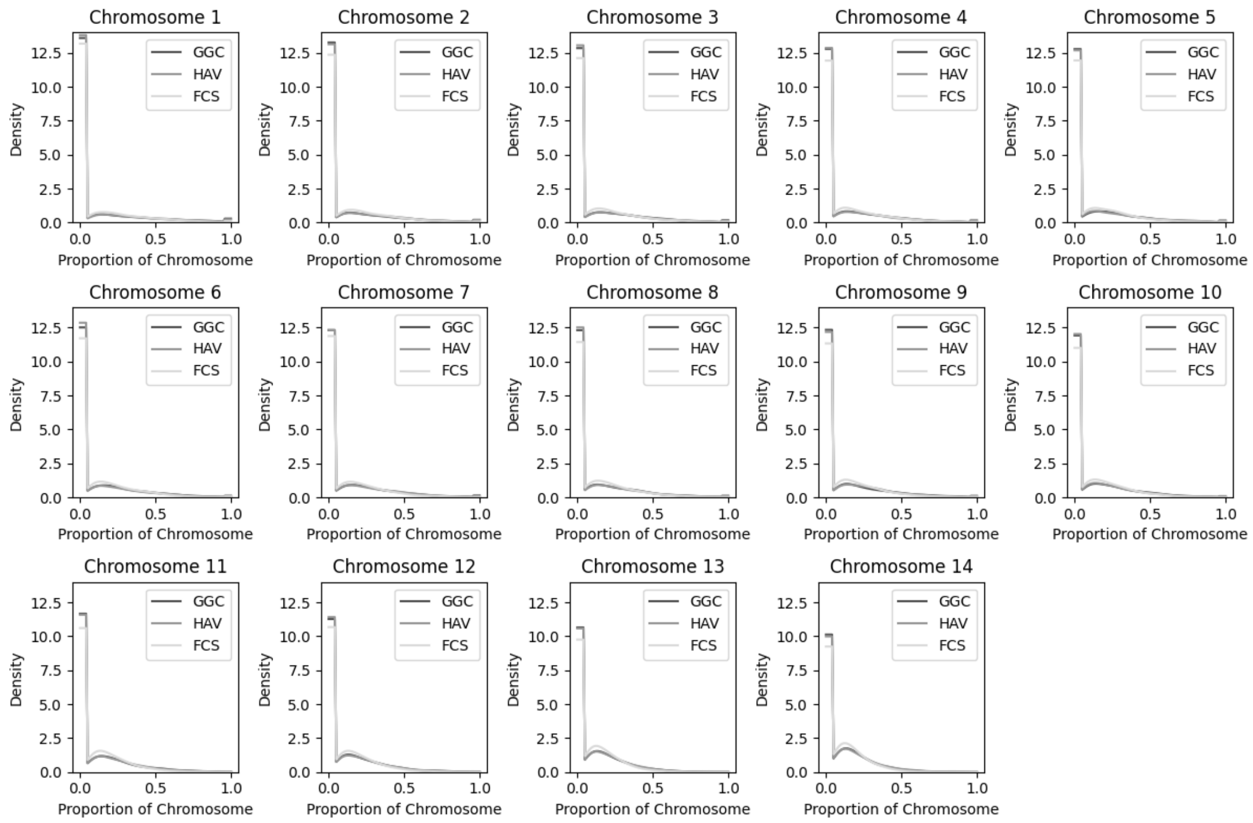


**Supplemental Figure 2C** Simulated max IBD segment block distribution for third-degree relatives.

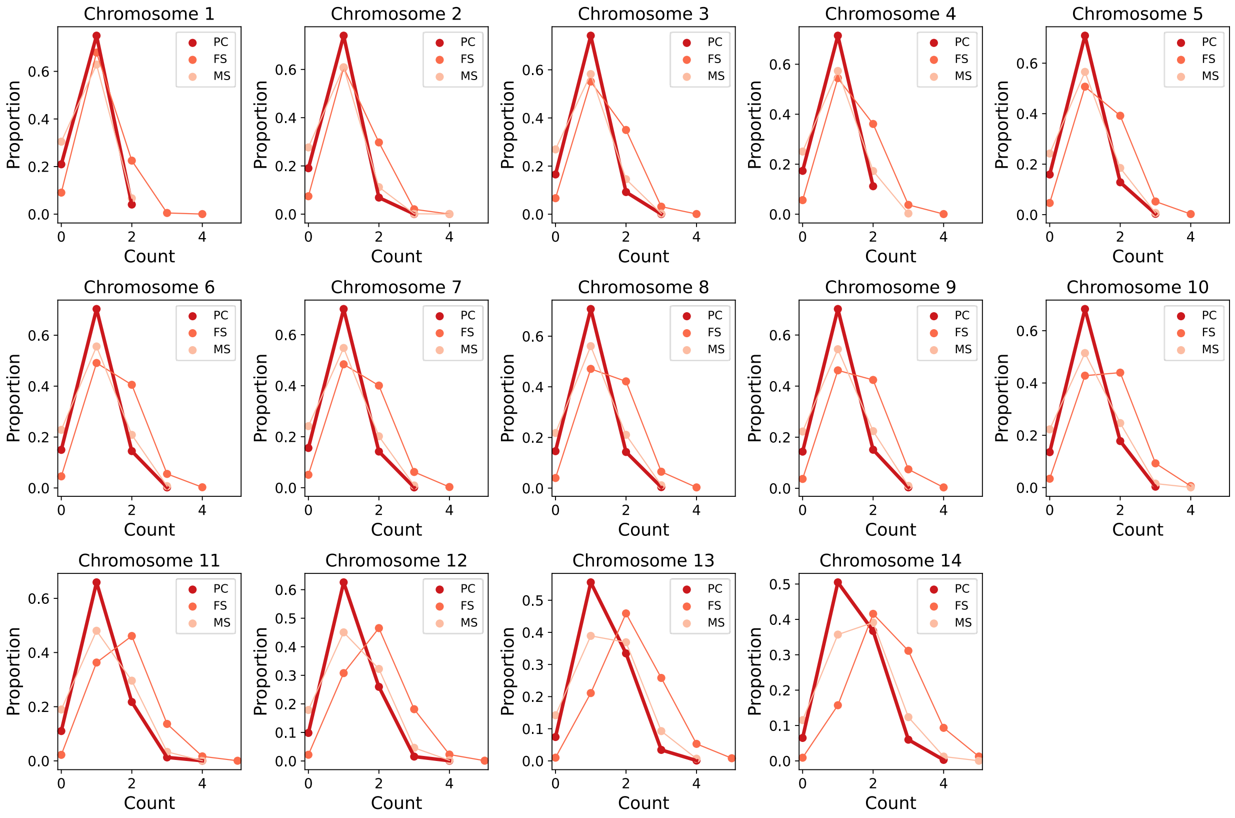


**Supplemental Figure 2D** Simulated max IBD segment count distribution for first-degree relatives.

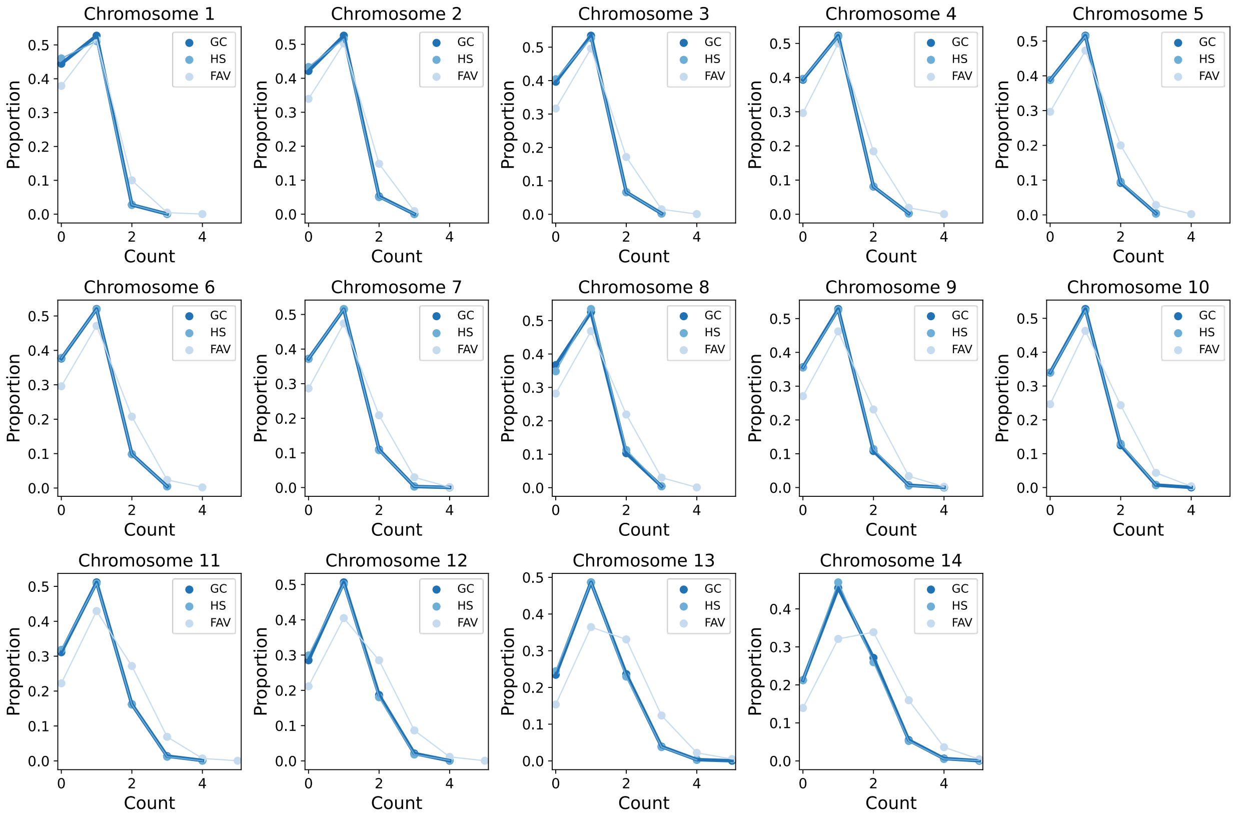

**Supplemental Figure 2E** Simulated max IBD segment count distribution for second-degree relatives.

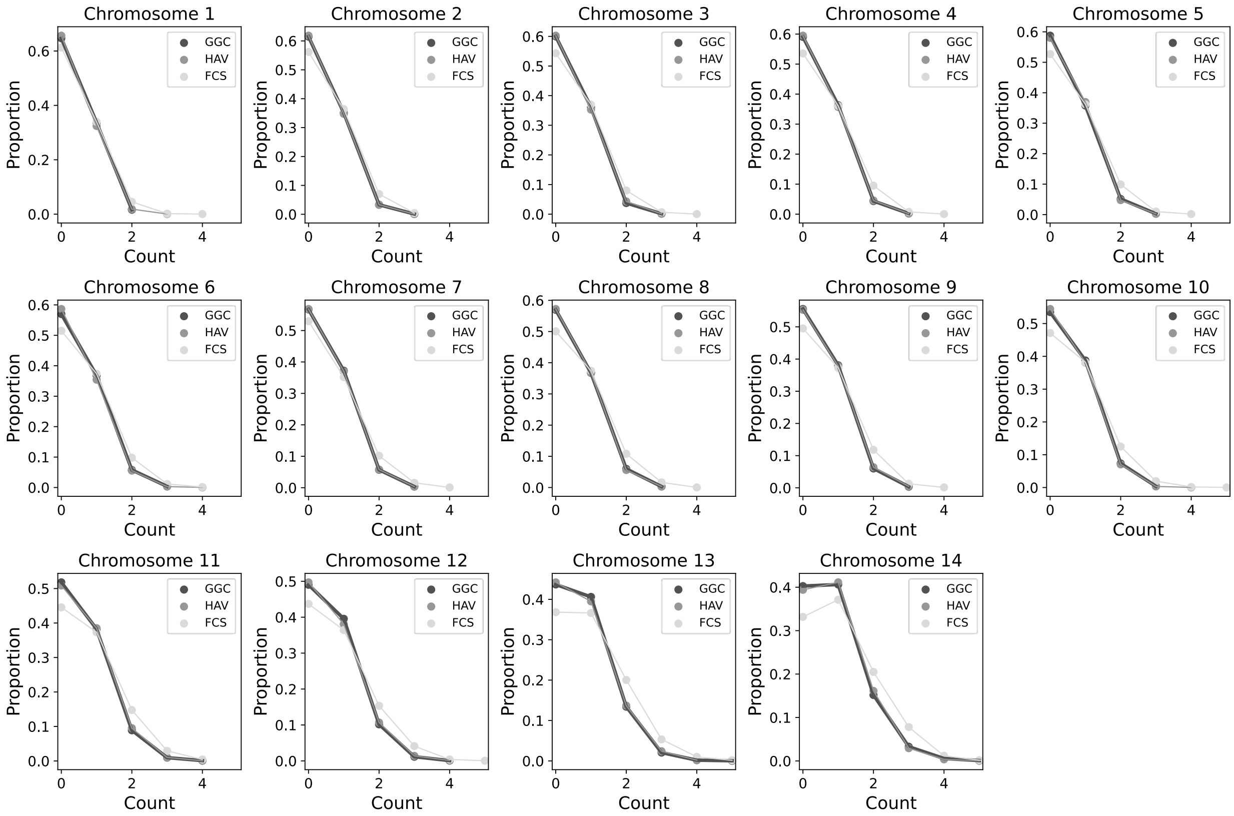


**Supplemental Figure 2G** Simulated max IBD segment count distribution for third-degree relatives.
